# Leveraging the effects of chloroquine on resistant malaria parasites for combination therapies

**DOI:** 10.1101/428284

**Authors:** Ana M. Untaroiu, Maureen A. Carey, Jennifer L. Guler, Jason A. Papin

**Affiliations:** Department of Biomedical Engineering, University of Virginia, Charlottesville, VA, USA; Department of Microbiology, Immunology, and Cancer Biology, University of Virginia, Charlottesville, VA, USA; Department of Biology, University of Virginia, Charlottesville, VA, USA

**Keywords:** Malaria, combination therapies, chloroquine, metabolic modeling

## Abstract

Malaria is a major global health problem, with the *Plasmodium falciparum* protozoan parasite causing the most severe form of the disease. Prevalence of drug-resistant *P. falciparum* highlights the need to understand the biology of resistance and to identify novel combination therapies that are effective against resistant parasites. Resistance has compromised the therapeutic use of many antimalarial drugs, including chloroquine, and limited our ability to treat malaria across the world. Fortunately, chloroquine resistance comes at a fitness cost to the parasite; this can be leveraged in developing combination therapies or to reinstate use of chloroquine. To understand biological changes induced by chloroquine treatment, we compared transcriptomics data from chloroquine-resistant parasites in the presence or absence of the drug. Using both linear models and a genome-scale metabolic network reconstruction of the parasite to interpret the expression data, we identified targetable pathways in resistant parasites. This study identified an increased importance of lipid synthesis, glutathione production/cycling, isoprenoids biosynthesis, and folate metabolism in response to chloroquine. We identified potential drug targets for chloroquine combination therapies. Significantly, our analysis suggests that the combination of chloroquine and sulfadoxine-pyrimethamine or fosmidomycin may be more effective against chloroquine-resistant parasites than either drug alone; further studies will explore the use of these drugs as chloroquine resistance blockers. Additional metabolic weaknesses were found in glutathione generation and lipid synthesis during chloroquine treatment. These processes could be targeted with novel inhibitors to reduce parasite growth and reduce the burden of malaria infections. Thus, we identified metabolic weaknesses of chloroquine-resistant parasites and propose targeted chloroquine combination therapies.

## Background

There are approximately 3.2 billion people at risk of malaria infection worldwide and the malaria parasites cause half a million deaths annually (1). Given the lack of a broadly effective vaccine, antimalarial drugs and protection from mosquito bites are essential in the control of malaria (2). The most lethal species of the protozoan parasite that causes malaria, *Plasmodium falciparum*, has acquired resistance to every antimalarial drug on the market (3,4). Since the development of novel antimalarials is slow, there is a need for combination therapies to target resistant parasites.

First introduced in 1934, chloroquine was a front-line antimalarial until the late 1950s when its heavy usage led to emergence of resistant *P. falciparum* strains near the Cambodia-Thailand border (5). Chloroquine resistance has now been confirmed in over 40 countries, making resistance to this drug a global concern (5). The mechanism of action of chloroquine is well studied. In intraerythrocytic trophozoite parasites, the drug blocks detoxification of heme, a byproduct of hemoglobin degradation (6). During the asexual intraerythrocytic-stages, the parasite imports host cell hemoglobin into its food vacuole (7,8). Proteases in the food vacuole degrade hemoglobin into free amino acids which are utilized in various growth processes (9). Heme is released during hemoglobin digestion and is essential for parasite growth as a cofactor for cytochromes in the parasite’s electron transport chain (10,11); however, elevated levels of intracellular heme can lead to cellular damage: oxidation of proteins, inhibition of proteases, and damage or lysis of membranes (12–14). Heme released from hemoglobin is detoxified through three known mechanisms: [1] polymerization into hemozoin crystals, [2] detoxification through interactions with hydrogen peroxide in the food vacuole, and [3] a glutathione-mediated degradation process in the cytoplasm (7,13,15,16). Chloroquine chemically binds to heme and the growing ends of hemozoin crystals, preventing crystallization-mediated heme detoxification (17,18).

Chloroquine-resistant parasites are able to export chloroquine (19), which reduces the accessibility of chloroquine to its heme and hemozoin targets (20,21). This resistance phenotype is mediated by mutations in the *P. falciparum* chloroquine resistance transporter *(pfcrt)* gene that results in the removal of drug from its functional site, the food vacuole (22,23). Numerous mutations are associated with chloroquine resistance, depending on the genetic background, and result in varying degrees of resistance (22,24–26). However, the substitution of lysine to threonine at position 76 in *pfcrt* is found in all *in vitro* chloroquine-resistant parasites (27,28).

Although chloroquine resistance is a well-studied case of antimalarial resistance, the mechanistic details remain poorly characterized. Studies on hemozoin formation estimate that only a third of heme released from hemoglobin is sequestered into hemozoin, suggesting a majority of heme is broken down through alternative, less well-characterized mechanisms, like peroxidative and glutathione degradation (16,29). Additional investigations are needed to understand the interplay between these mechanisms, which may become important in situations where resistant parasites are exposed to chloroquine. This situation is a greater possibility today, as some malaria endemic countries consider reintroducing chloroquine into their treatment regimens. Chloroquine resistance confers a fitness cost; thus, resistance alleles do not become fixed and decline in prevalence after chloroquine use is discontinued (30,31). As a result, clinical trials have confirmed that reintroduction of chloroquine is highly efficacious (32,33). Based on these findings, a reintroduction of chloroquine as a part of combination therapies may be an effective treatment strategy.

This study investigates the effects of chloroquine on chloroquine-resistant parasites to identify partner drugs that could be used in combination therapies to accelerate the reintroduction of chloroquine. We integrated the transcriptomics data from chloroquine-resistant parasites with or without chloroquine treatment (34) into a *P. falciparum* genome-scale metabolic network reconstruction to predict large-scale metabolic changes initiated by drug treatment, as we previously performed to study artemisinin resistance (35). We identified shifts in metabolic flux and clinically available inhibitors that may partner well with chloroquine to target chloroquine resistant parasites.

## Methods

### Differential Gene Analysis

Expression data from *in vitro* K1 parasites untreated or treated with EC_50_ concentrations of chloroquine for 4hr and 24hrs were used to investigate the transcriptional effects of chloroquine treatment (normalized expression data obtained via GEO accession number: GSE31109) (34). Exposure for 4 and 24 hours were defined as short and long-term treatment, respectively. Probes with missing expression values were set equal to the lowest expression value across all replicates for that probe (35). Mean or median values across replicates were not used to replace missing expression values to avoid skewing the data with outliers. Genes with multiple probes were filtered from the data to ensure each gene corresponds to only one microarray probe. For repeating probes, only expression data from the probe with the largest variance across all replicates were retained. Means from repeating probes were not used to prevent averaging true signal with background noise. Probes with missing gene IDs were removed from the dataset.

The R package limma was used to conduct differential gene expression analysis **(Figure 1)**. Limma uses linear models to fit a data set and empirical Bayes statistical methods to determine variability and log-fold changes in gene expression (36). Significance values from this analysis (p-values) were modified using false discovery rate (FDR) correction in order to control for the parallel manner of comparing genes using limma, instead of testing genes in isolation. Differentially expressed genes were characterized as genes having a fold change (FC) greater than 2 or less than 0.5 and a FDR-adjusted p-value less than 0.05.

**Figure 1:**
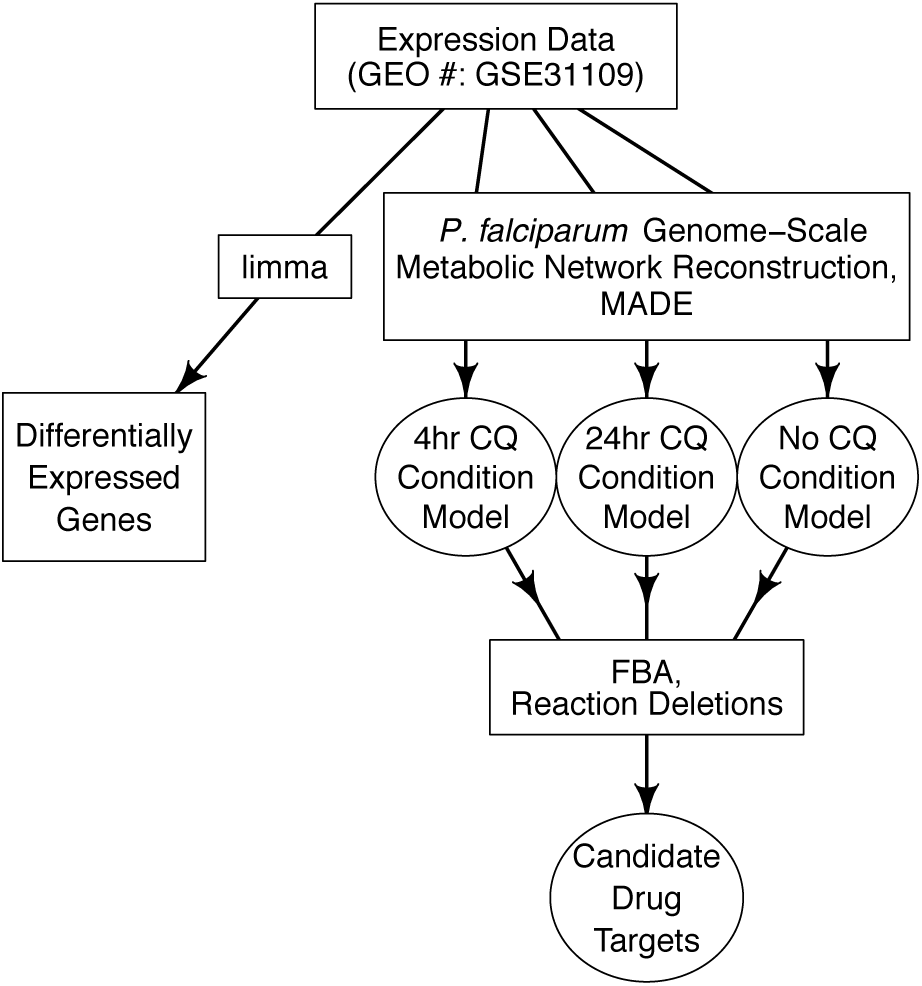
Overview of Computational Approach. The R package limma was used to find differentially expressed genes (left square). Metabolic Adjustment for Differential Expression (MADE) algorithm was used to produce the condition-specific models (right 3 circles). Flux balance analysis (FBA) and simulated reaction deletions predicted drug targets for chloroquine-treated parasites. CQ = Chloroquine.

### Model Curation

Additional reactions were included into the *P. falciparum* model based on supporting experimental evidence (35). More specifically, hydrogen peroxide was incorporated as a byproduct of hemoglobin digestion based on studies of the chemical steps in hemoglobin breakdown (37). Reactions for heme degradation via glutathione and hydrogen peroxide were incorporated based on supporting literature (38,39). See **Supplemental Table 1** for modified reactions.

### Condition-specific Model Generation

Condition-specific models were produced by integrating gene IDs with corresponding log FCs and FDR-adjusted p-values from the 4 and 24 hour treatment conditions into a curated version (see above) of the intraerythrocytic-stage *P. falciparum* genome-scale metabolic network reconstruction (35) (**Figure 1**). Data integration was conducted by using the Metabolic Adjustment for Differential Expression (MADE) algorithm; this algorithm accounts for the expression state of a gene with a weighted consideration of the statistical significance of the gene expression changes (40). Condition-specific models were generated for each condition (4hr chloroquine-treated, 24hr chloroquine-treated, and untreated) using varying growth thresholds (30 – 80%). This threshold represents the fraction of metabolic objective required for the output condition-specific model. We explored the effects of this threshold on condition-specific models to find minor differences in results. This paper reports common results across all models.

### Reaction Essentiality Predictions and Flux Analysis

Essential reactions for each condition-specific model were generated by sequentially removing single reactions and testing the modified model for growth using flux balance analysis (41) (**Figure 1**). Reactions were delineated as essential when the removal of the reaction from the model resulted in a decrease of the growth rate by 60% or more compared to the untreated model. This growth rate threshold was chosen to avoid a too conservative definition of essentiality, but to also ensure a significant growth defect. Existing algorithms in the COBRA toolbox were used to calculate ranges of possible flux levels through reactions in the condition-specific models (42,43). Generation of condition-specific models, reaction essentiality, and flux analysis were conducted using MATLAB vR2014a, COBRA Toolbox v3.0, and Gurobi 6.5.2 solver.

### Enrichment Analysis

The enrichment of metabolic subsystems among essential reactions were investigated. The purpose of this analysis is to determine whether a certain subsystem appears to a greater or lesser extent than expected by chance. Corresponding metabolic subsystems for genes and reactions were derived from the genome-scale metabolic model used in this study (35). We tested for enrichment of subsystems among essential genes with reference to the unconstrained model using a Fisher’s exact test. Significant subsystems were defined as having a FDR-adjusted p-value less than 0.05. RStudio v3.3.0 was used for the differential gene and enrichment analysis.

## Results

### Resistant parasites respond to chloroquine pressure with moderate but statistically significant gene down-regulation

Since this analysis was not performed in the original study (34), differentially expressed genes (DEGs) were identified for both short and long-term treatment relative to untreated trophozoite expression. The majority of genes do not show a significant change in expression in response to chloroquine treatment (genes represented by points in black, **Figure 2**). Out of a total of 5,121 genes, there were 97 and 166 differentially expressed genes (DEGs) in short- and long-term treatment conditions, respectively (represented by points in red, **Figure 2**). In both conditions, more genes were significantly down-regulated than up-regulated in response to chloroquine treatment (95.9% for short-term and 83.1% for long-term treatment, fold change less than 0.5, **Suppl. Table 2**). Up-regulated genes showed very moderate over-expression (fold change between 2 and 2.5) and included one metabolic gene (**Suppl. Table 3**). No evidence was found to suggest these DEGs are directly involved in chloroquine treatment or resistance.

**Figure 2:**
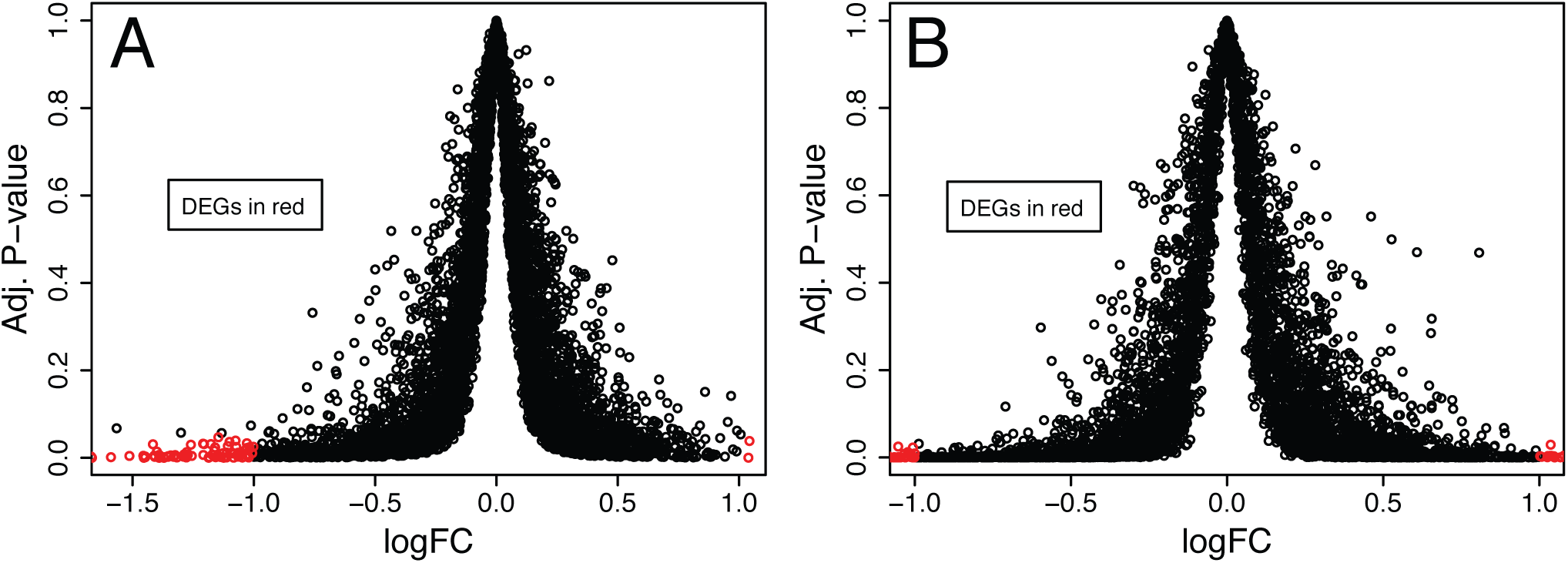
Resistant Parasites Respond to Drug by Downregulating Expression. FDR-adjusted p-values and fold changes (FC, reported as log values) for genes during (A) short-term treatment and (B) long-term treatment with chloroquine. Adjusted p-values represent the significance level of changes in expression. Fold change quantifies the variation in the gene expression relative to untreated resistant parasite expression. DEGs = differentially expressed genes.

### Identifying metabolic weakness of chloroquine-treated resistant parasites using metabolic modeling

In order to understand the system-wide context for these gene expression changes, we integrated transcriptomics data into a genome-scale metabolic network reconstruction of intraerythrocytic-stage *P. falciparum* to generate three condition-specific models (untreated, short- and long-term chloroquine treatment). We then predicted reactions essential for parasite growth. One hundred and sixty-four metabolic reactions (out of 1197) are essential in all three models (**Figure 3A**, center); these reactions represent core metabolic pathways of the parasite.

**Figure 3.**
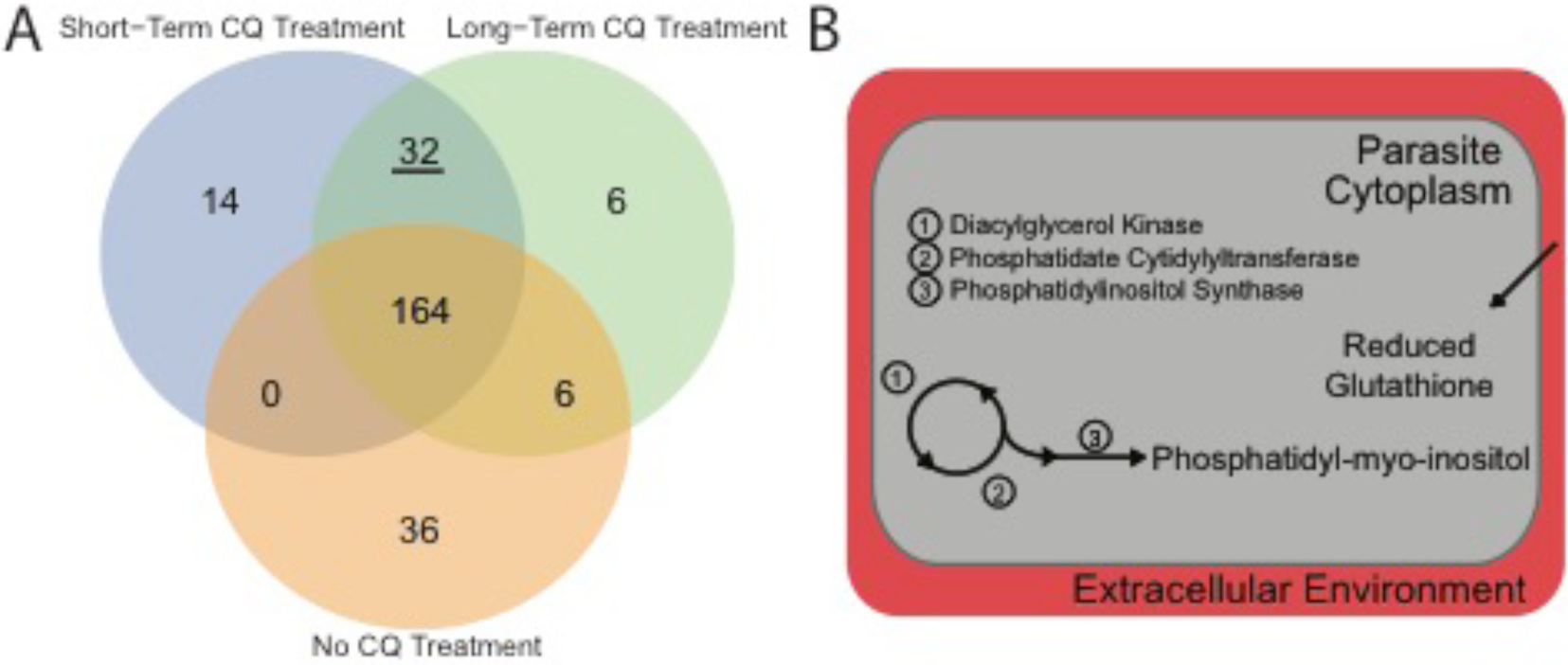
Chloroquine-treated parasites have new metabolic weaknesses. (**A**) Comparison of essential reactions in the three condition-specific models (one for each condition: short-term and long-term chloroquine treatment, as well as the untreated condition). (**B**) Illustration of common essentiality predictions (underlined in A) between the drug-treatment models, including inositol phosphate metabolism and glutathione import are represented. These enzymes could be targeted in these resistant parasites in combination with chloroquine; resultant combination therapies would specifically target resistant parasites during chloroquine treatment, not wild-type parasites or single-drug treatment. Red region depicts the host red blood cell and grey is the parasite’s cytoplasm.

Reactions that become essential during chloroquine treatment highlight weak points in the metabolic network and can be exploited as drug targets in chloroquine-resistant parasites. We found 210 and 208 essential metabolic reactions in short- and long-term treatment, respectively. Thirty-two were shared between the two conditions **(Suppl. Table 5, Figure 3**); fifteen of these are involved in phosphatidylethanolamine and phosphatidylserine metabolism, seven in phospholipid utilization, and one in the import of reduced glutathione (examples summarized in **Figure 3B**; **Suppl. Table 5**). When we performed subsystem enrichment on the essential reactions, the following subsystems were over-represented: phosphatidylethanolamine and phosphatidylserine metabolism (p-value = 4.84E-6), tRNA synthesis (p-value = 0.024), and phospholipid utilization (p-value = 0.037). Reactions involved in transport and lipid metabolism were under-represented in chloroquine treated models (p-value = 0.0061 and 2.51E-8, respectively).

Fourteen reactions are essential in only the short-term treatment model **(Figure 3A** in blue, **Suppl. Table 6**). Of these essential reactions, five reactions are involved in the *de novo* synthesis of thiamine diphosphate, the active form of vitamin B1 **(Figure 4** in white). Eight reactions were uniquely essential in the long-term treatment model **(Figure 3A** in green, **Suppl. Table 7)**, notably the conversion of chorismate into 4-aminobenzoate for folate metabolism is essential **(Figure 5** in black).

**Figure 4.**
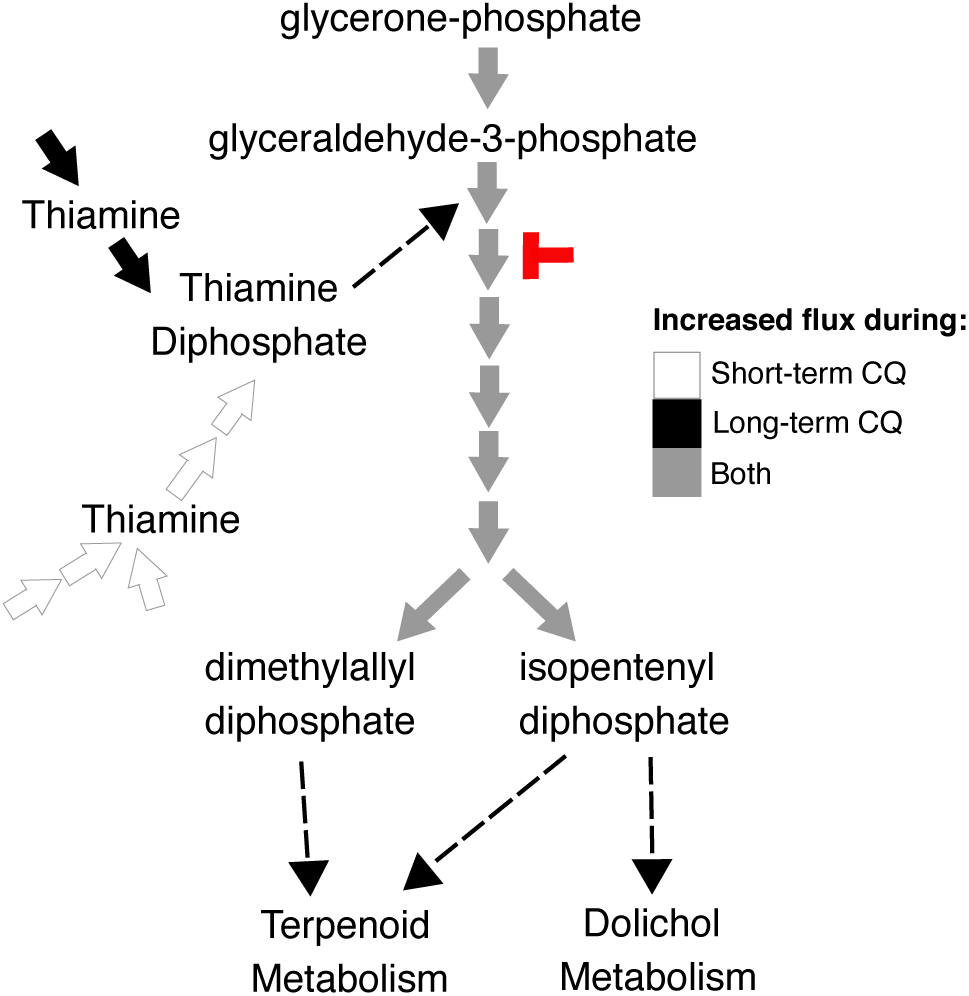
Increased Flux in Isoprenoids Metabolism in Response to Chloroquine. Illustration of isoprenoid metabolism reactions showing increases in flux levels in both chloroquine-treated models versus untreated models (represented in grey). 1-deoxy-D-xylulose-5-phosphate synthase utilizes thiamine diphosphate as a cofactor (marked by dotted arrow). We also predict a shift in thiamine diphosphate production, where the *de novo* synthesis pathway (shown in white) and the import pathway (shown in solid black) is used during short- and long-term treatment, respectively. Pharmacologic inhibitors target isoprenoid metabolism (fosmidomycin, indicated in red). Connectivity to other metabolic pathways are shown (dotted lines).

**Figure 5.**
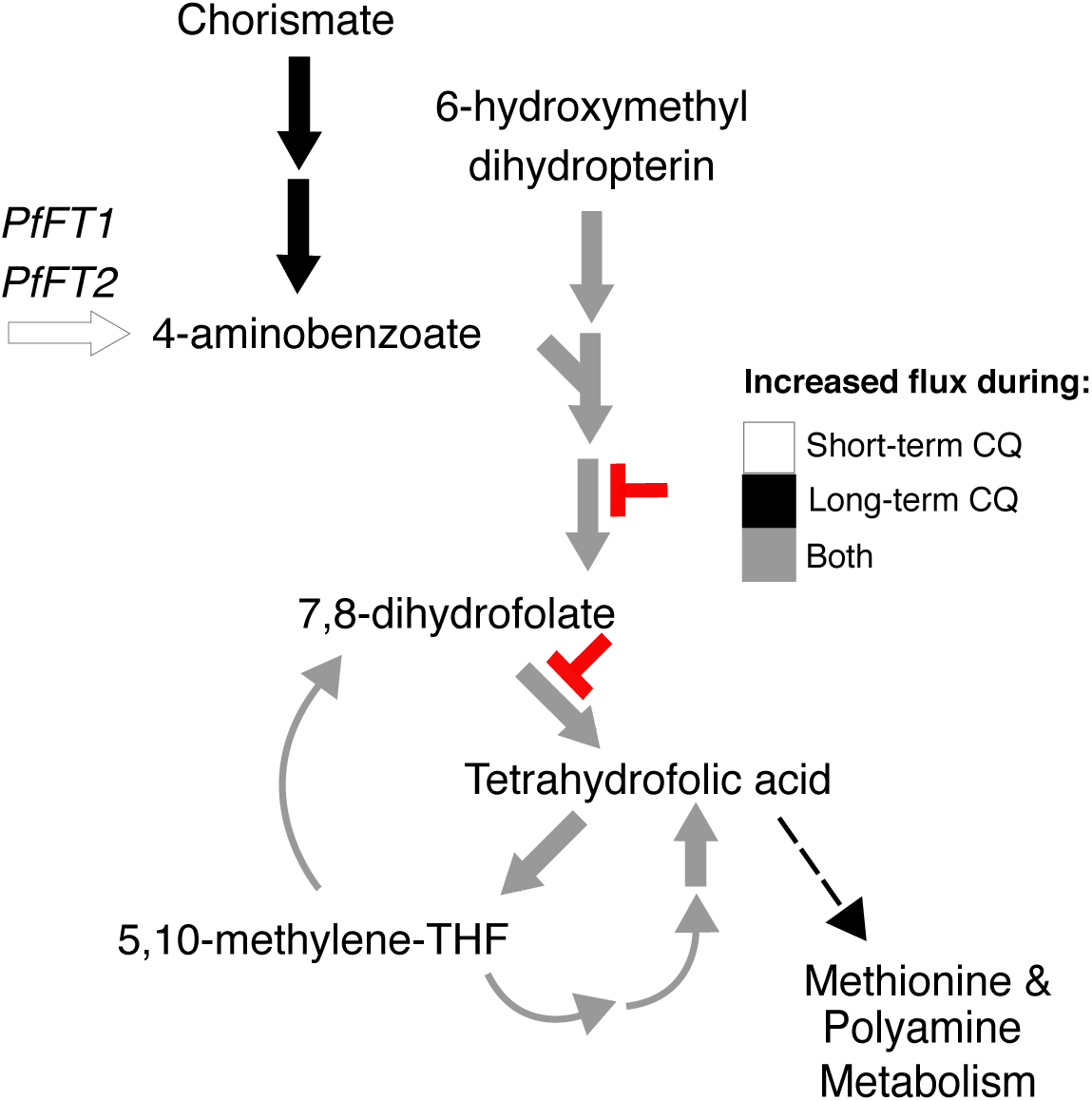
Increased Flux in Folate Metabolism in Response to Chloroquine. Illustration of folate metabolism reactions showing increases in flux levels in both chloroquine-treated models versus untreated models. We predict a shift in 4-aminobenzoate acquisition between short- and long-term chloroquine treatment, and elevated flux through all downstream steps of folate metabolism. Clinically available drugs target this pathway (sulfadoxine and pyrimethamine, indicated in red). Both enzymes targeted by these antimalarials (dihydropteroate synthase and dihydrofolate reductase) carry increased flux in both models of short- and long-term chloroquine treatment. Connectivity to other metabolic pathways are shown (dotted lines).

We explored why thiamine diphosphate synthesis was essential only in the short-term treatment model. *P. falciparum* has an alternate route for thiamine diphosphate production (termed ‘import pathway’, **Figure 4**), where thiamine is imported into the parasite’s cytoplasm and phosphorylated by thiamine diphosphokinase to form thiamine diphosphate. For the untreated and long-term treatment models, only the import pathway is active. For the short-term treatment model, only the *de novo* synthesis pathway has activity. These results suggest a switch in thiamine diphosphate production in early stages of drug treatment.

Six enzymes are known to utilize thiamine diphosphate (also called thiamine pyrophosphate) as a cofactor in *P. falciparum* (44–48). Flux levels of these reactions predicted from flux balance analysis were investigated to understand thiamine diphosphate usage and essentiality. Flux balance analysis simulates steady-state flux values through the network’s reactions and predicts the reactions needed to maximize the objective function, which was selected as the biomass equation; thus, this analysis predicts reactions needed for growth. Four of these thiamine diphosphate dependent enzymes (pyruvate dehydrogenase, 2-oxoglutarate dehydrogenase, 3-methyl-2-oxobutanoate dehydrogenase, and branched-chain-alpha-keto-acid dehydrogenase) are predicted to carry no flux in treated and untreated parasites. In response to drug treatment, flux changes through two thiamine diphosphate-dependent reactions: flux through the reaction catalyzed by 1-deoxy-D-xylulose phosphate synthase increases (in isoprenoids metabolism) and flux through the reaction catalyzed by transketolase decreases (in pentose phosphate pathway). Flux of the other reactions in isoprenoids metabolism are also consistently greater in response to chloroquine treatment **(Figure 4** in grey**; Suppl. Table 8)**. Compared to untreated models, short- and long-term treatment show a 62 – 83% and 24% increase in flux, respectively.

To next focus on the reactions essential in only the long-term treatment model, we explored the utilization of chorismate, a precursor of folate. During long-term chloroquine treatment, synthesis of 4-aminobenzoate from chorismate is used, rather than importing 4-aminobenzoate from the host cell **(Figure 5)**. Thus, the conversion of chorismate into 4-aminobenzoate is essential. In contrast, short-term chloroquine-treated parasites predominately rely on membrane folate transporters (PfFT1 and PfFT2) for 4-aminobenzoate **(Figure 5)**. Downstream steps in folate metabolism, including dihydropteroate synthase and dihydrofolate reductase, carry a 62.4% and 24.2% increase in flux during short- and long-term chloroquine treatment, respectively **(Figure 5** in grey; **Suppl. Table 9**).

## Discussion

In this study, we used transcriptomics data to investigate metabolic shifts resulting from chloroquine treatment of resistant parasites. We performed differential expression analysis and integrated expression data into a genome-scale metabolic model to identify targetable weaknesses of chloroquine-treated resistant parasites. Using this approach, we identified metabolic pathways that, if targeted, could be developed as partner drugs for chloroquine combination therapies.

### Chloroquine affects resistant parasites

Importantly, we observed a significant down-regulation of genes in response to chloroquine treatment (**Figure 2**). This suggests that chloroquine treatment continues to affect the resistant parasite, despite the parasite’s resistance to the drug’s cytocidal effects. This observation is consistent with growth defects observed in chloroquine-treated resistant parasites (20% and 45% inhibition after 8 and 48 hours, respectively (49)). Moreover, the effect of chloroquine is widespread and not just a function of a single gene, as evident by the numerous differentially expressed genes, and the multiple nonspecific resistance alleles (50,51). Longer exposure time exacerbates this effect, as illustrated by the increase in DEGs with the 24-hour drug treatment (**Figure 2**).

### Novel proposed targets against chloroquine-resistant parasites

We propose that resistant parasites are using different metabolic pathways when in the presence of chloroquine because there are treatment-associated essential reactions. Subsystem enrichment of essentiality predictions suggests many shifts in lipid metabolism. The increased importance of phospholipids in both treatment conditions (Figure 3B) may represent the parasite’s attempt to counteract the effects of chloroquine treatment. This response could occur through a number of routes: first, additional lipid species may be required to repair cellular membranes damaged by the build-up of intracellular heme during chloroquine treatment (14). Second, lipids themselves have been shown to contribute to the detoxification of heme into hemozoin (52,53). The demand for lipids for these roles may represent a targetable metabolic weakness of drug-treated chloroquine-resistant parasites.

The transport of glutathione from the extracellular environment into the cytoplasm is predicted to be essential during chloroquine treatment **(Figure 3B)**. Glutathione is involved in the degradation of non-polymerized heme, in addition to being involved in managing oxidative stress in the parasite (15). Since these reactions are only essential during chloroquine treatment, this result suggests the activation of these reactions may be a direct result of drug pressure placed on the parasite and the accompanying cellular damage. This result is supported by observed correlations between chloroquine resistance and intracellular glutathione levels (54–56). The competitive inhibition of glutathione degradation by heme also supports the increased importance of glutathione accumulations to counteract chloroquine pressures (57). Thus, glutathione is essential in combating the effects of chloroquine and can be considered another metabolic weakness of resistant parasites.

### Partner drugs for chloroquine combination therapies

Folate metabolism is needed for DNA synthesis and metabolism of certain amino acids (52). Interestingly, downstream steps in folate metabolism, including dihydropteroate synthase and dihydrofolate reductase, are predicted to carry more flux during chloroquine treatment **(Figure 5)**, implying they are necessary for survival or tend to be overexpressed during treatment. This result suggests that this pathway has increased importance under chloroquine treatment and could be targeted in combination therapies. Recent clinical use of such a combination therapy supports this conclusion; chloroquine in combination with inhibitors of dihydrofolate reductase and dihydropteroate synthase (sulfadoxine-pyrimethamine) is effective against chloroquine-resistant parasites (58–60). Our results suggest that chloroquine-resistant parasites are more susceptible to these drugs than sensitive parasites and our modeling approach provides a mechanistic explanation.

Unique to our study, we predict a novel role for isoprenoids synthesis in chloroquine-resistant parasites. Under chloroquine treatment, there is increased flux through reactions in the non-mevalonate pathway for isoprenoids metabolism (**Figure 4**), the only synthesis pathway for isoprenoids in *P. falciparum* (61,62). This pathway is thiamine diphosphate-dependent and we also observed a switch in thiamine scavenging to *de novo* synthesis (**Figure 4**), highlighting the dynamic state of these pathways. Our computational analysis suggests that chloroquine-resistant parasites have increased susceptibility to non-mevalonate pathway inhibitors, such as fosmidomycin and its derivative, FR-900098 **(Figure 4)** (63). This metabolic weakness represents an ideal target since the non-mevalonate pathway is constitutively essential, with increased usage in chloroquine-treated resistant parasites (64). Fosmidomycin alone is moderately effective against chloroquine-resistant parasites (65) and there is no additivity between fosmidomycin and chloroquine *in vitro* in a pool of chloroquine-sensitive and resistant parasites (66). Thus, we propose that these parasites are even more susceptible to fosmidomycin while under chloroquine treatment. Thus, we have identified a novel combination therapy involving readily available antimalarials that may inhibit the growth of chloroquine-resistant parasites.

## Conclusions

The fitness cost associated with chloroquine resistance means that once drug pressure is removed, sensitive parasites again become prominent in the population. Due to its low cost and easy access in malaria endemic countries, chloroquine use is being considered in areas where the return to sensitivity has been confirmed. However, we must be deliberate about how this drug is reinstated to avoid the rapid return of resistant parasites. Thus, we identified potential drug targets for chloroquine combination therapies, using metabolic modeling of resistant parasites under chloroquine treatment. Significantly, we identified that the combination of chloroquine with sulfadoxine-pyrimethamine or fosmidomycin may be more effective against chloroquine-resistant parasites than either drug alone; further studies will explore the use of these drugs as chloroquine resistance blockers. Additional metabolic weaknesses were found in glutathione generation and lipid synthesis during chloroquine treatment. These processes could be targeted with novel inhibitors to reduce parasite growth, thus reducing the burden of malaria infections.

## Supplemental Information

**Suppl. Table 1:**
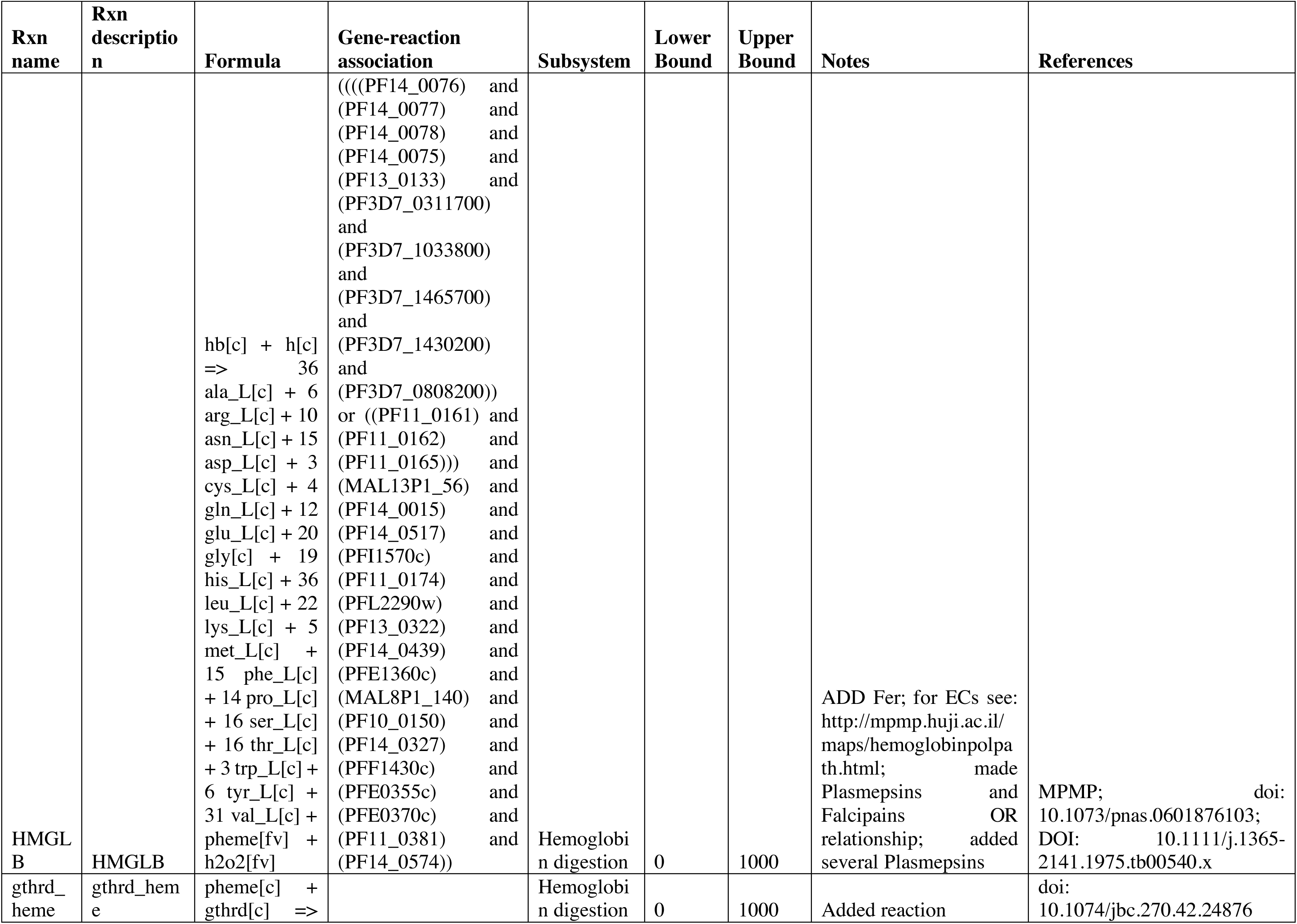
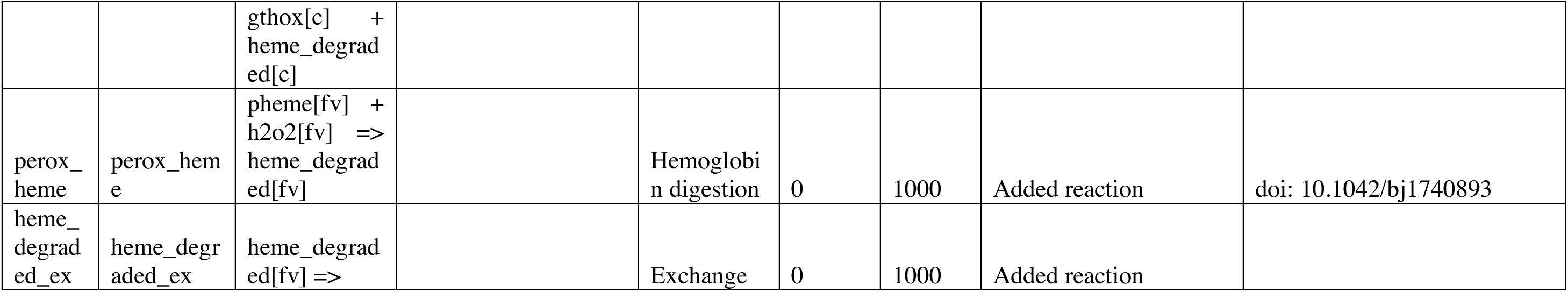
Reactions modified to curate the *Plasmodium falciparum* genome-scale metabolic model.

**Suppl. Table 2:**
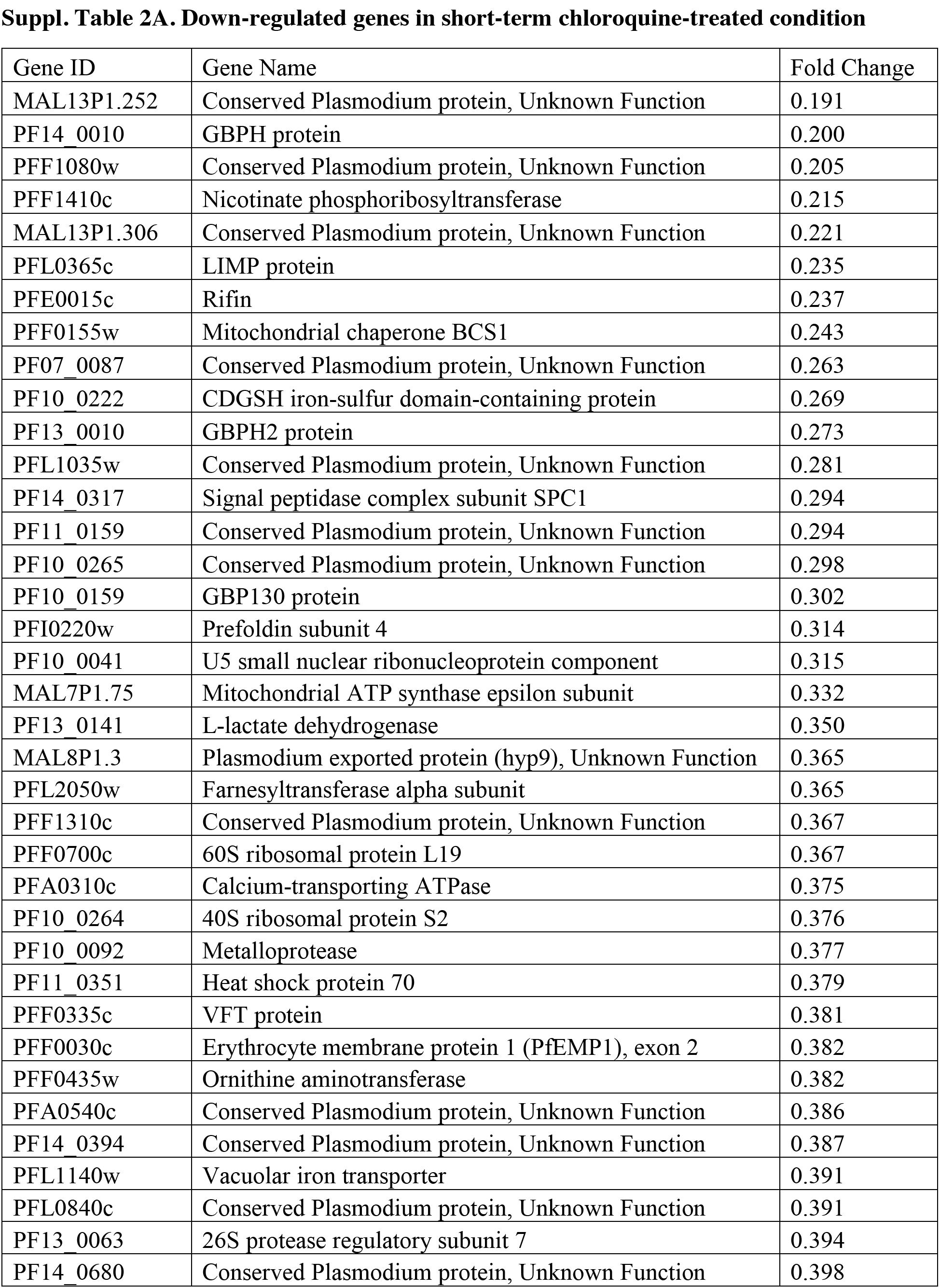
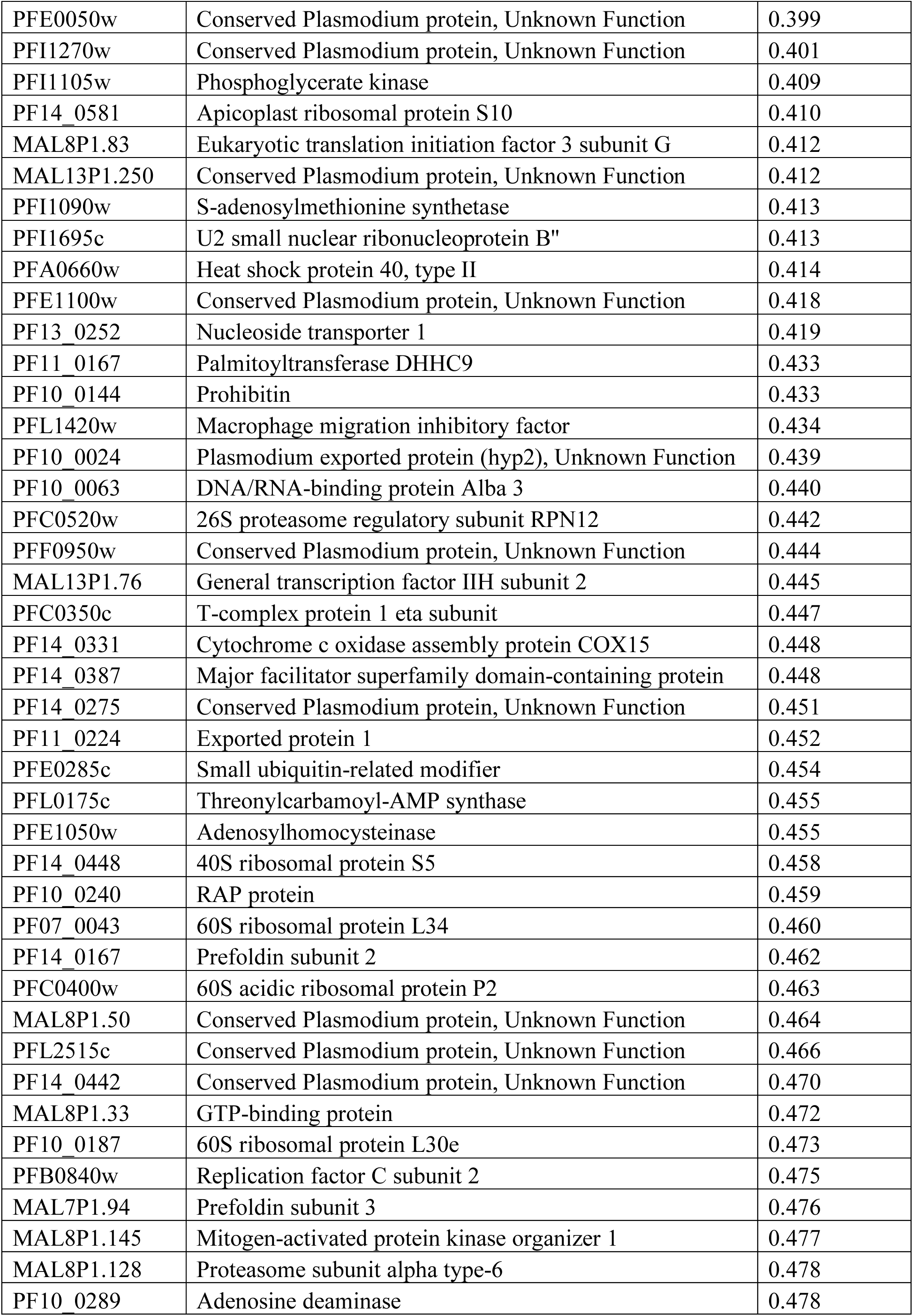
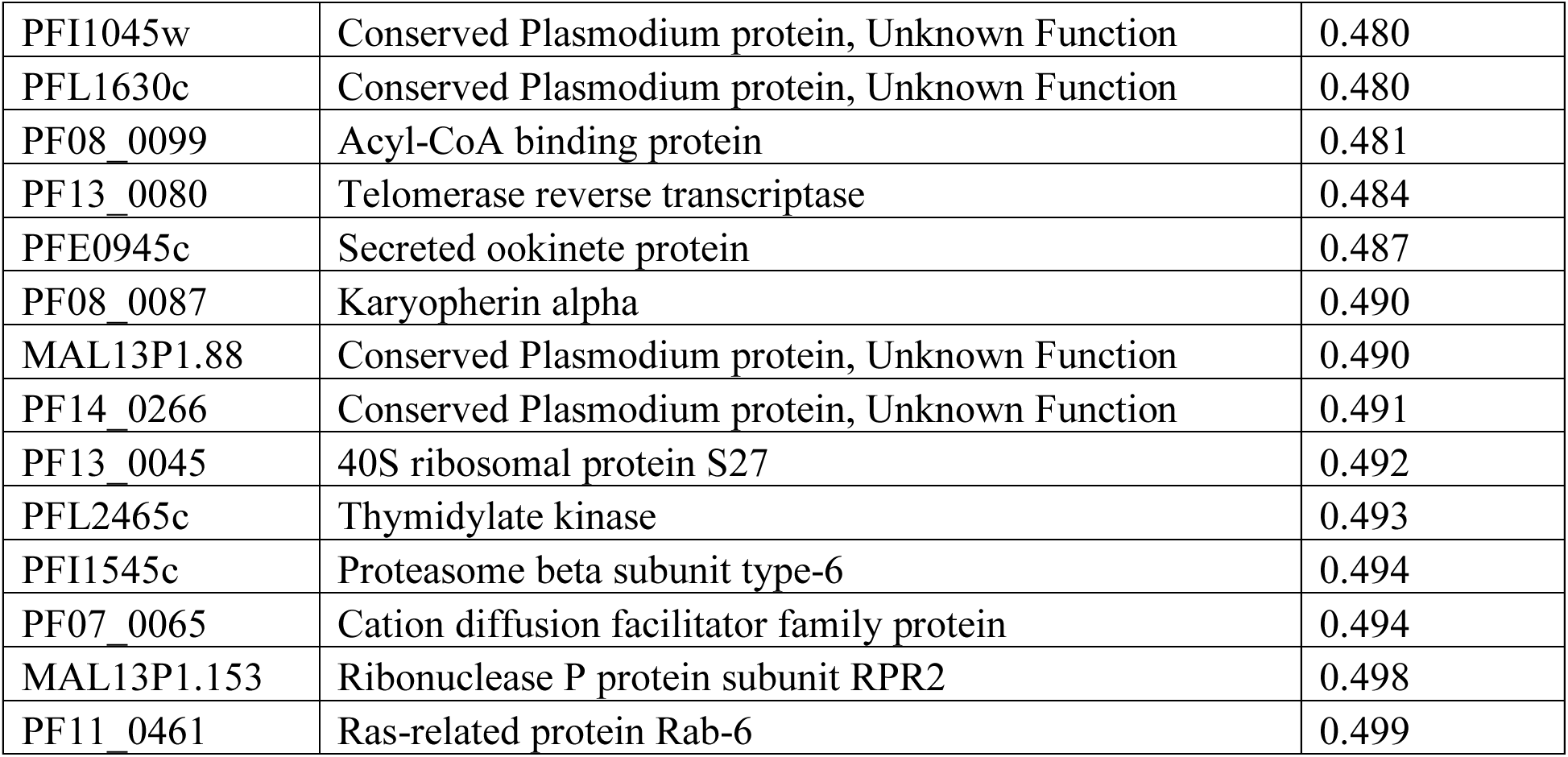
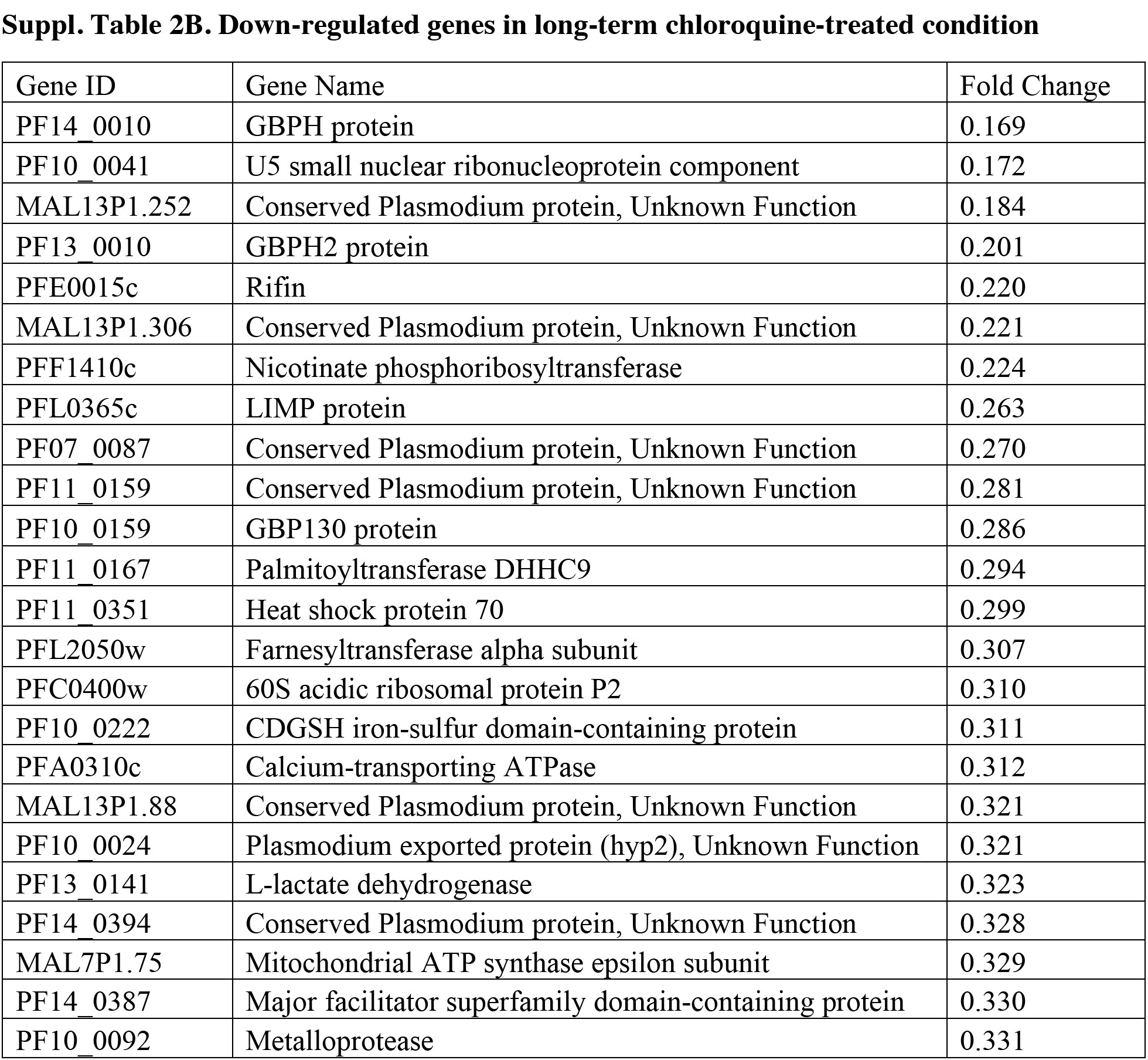
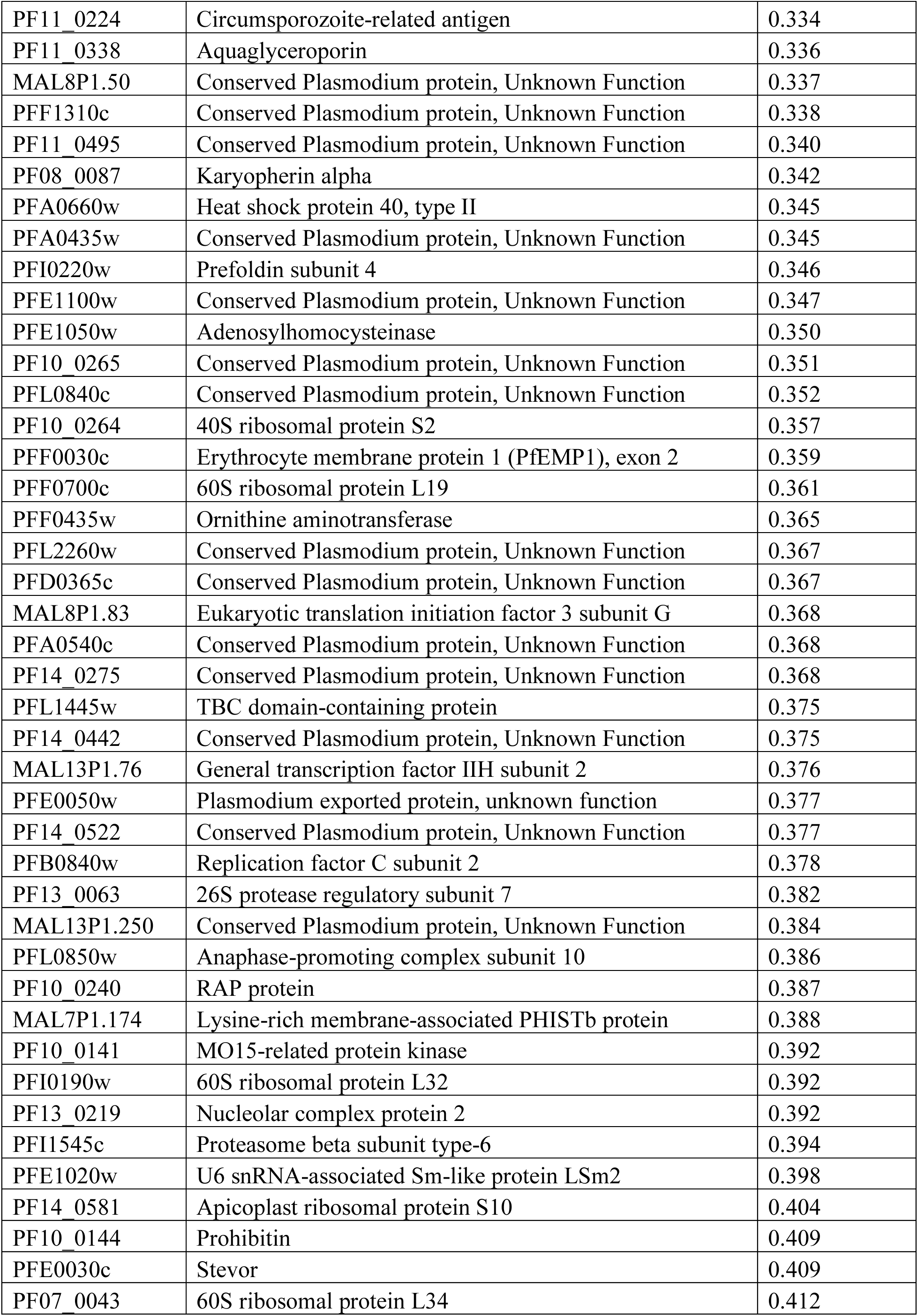
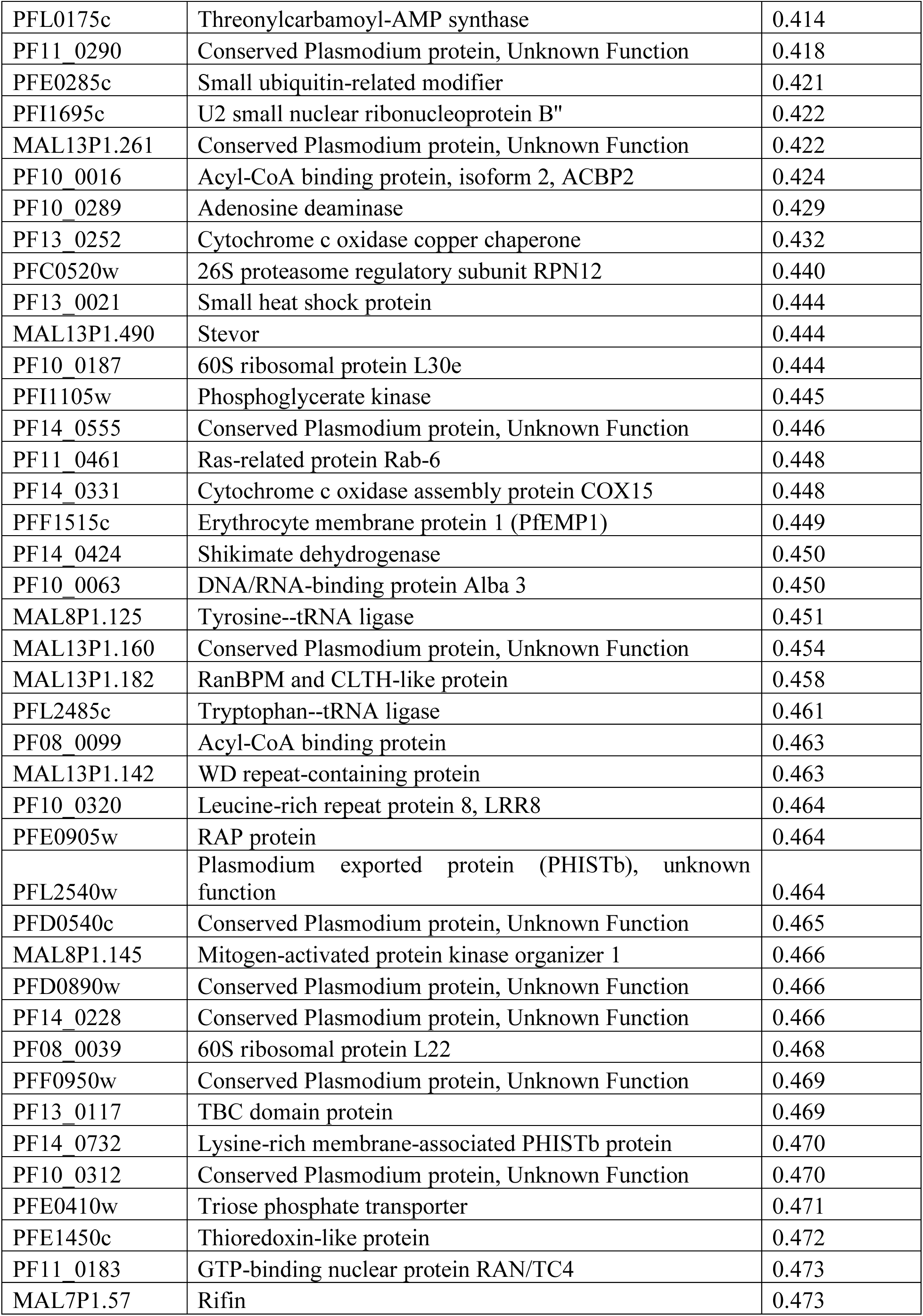
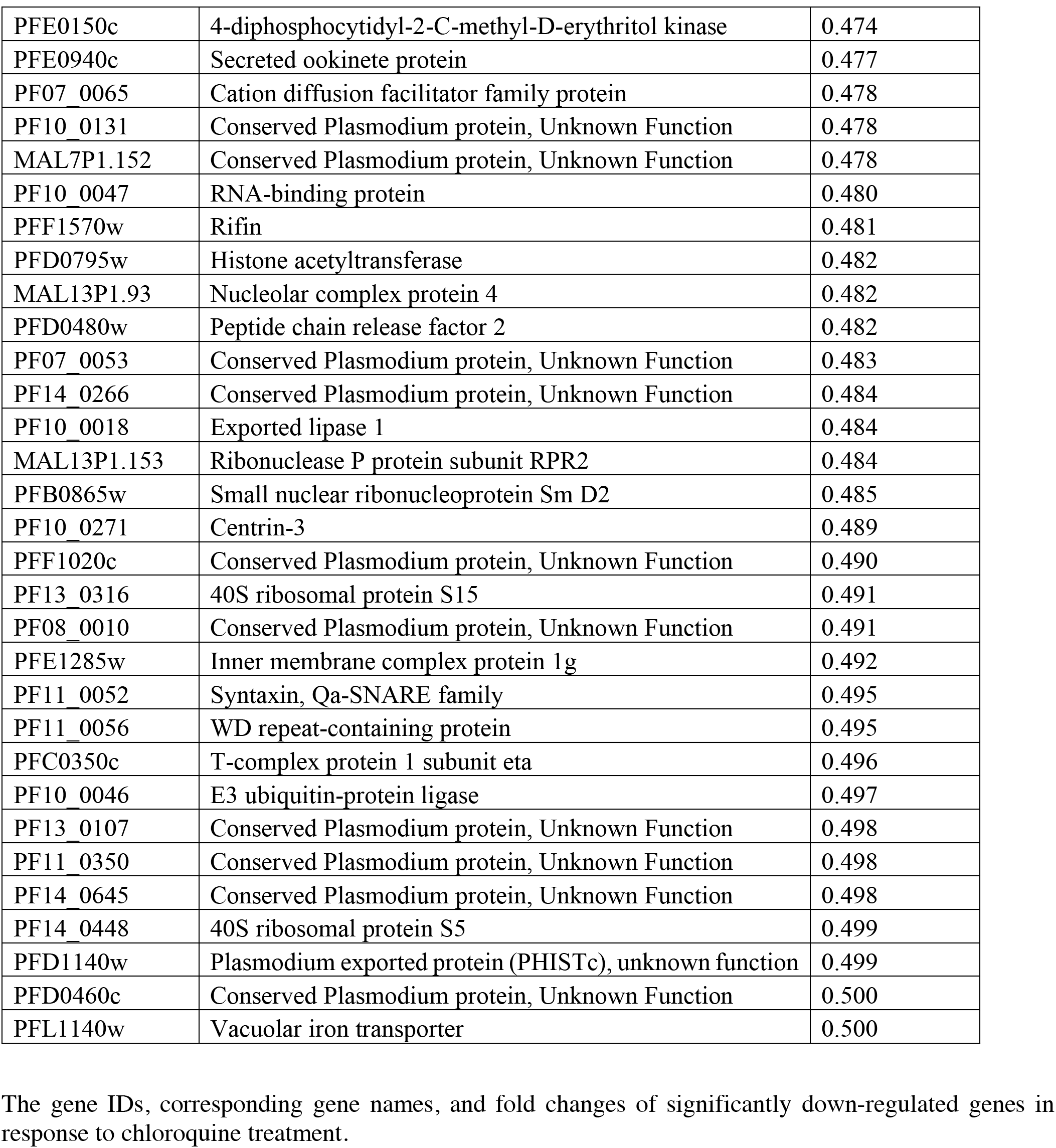
Down-regulated Differentially Expressed Genes in (A) Short and (B) Long-term Chloroquine-treated Parasites.

**Suppl. Table 3:**
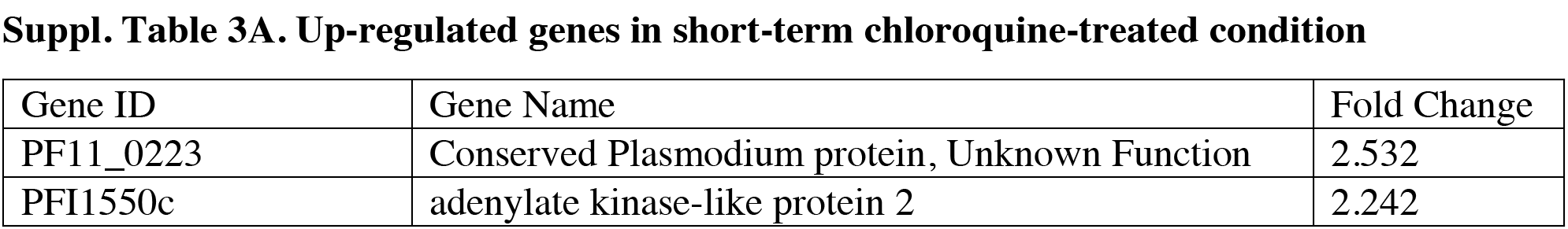
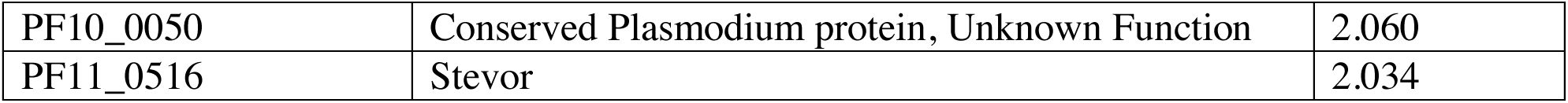
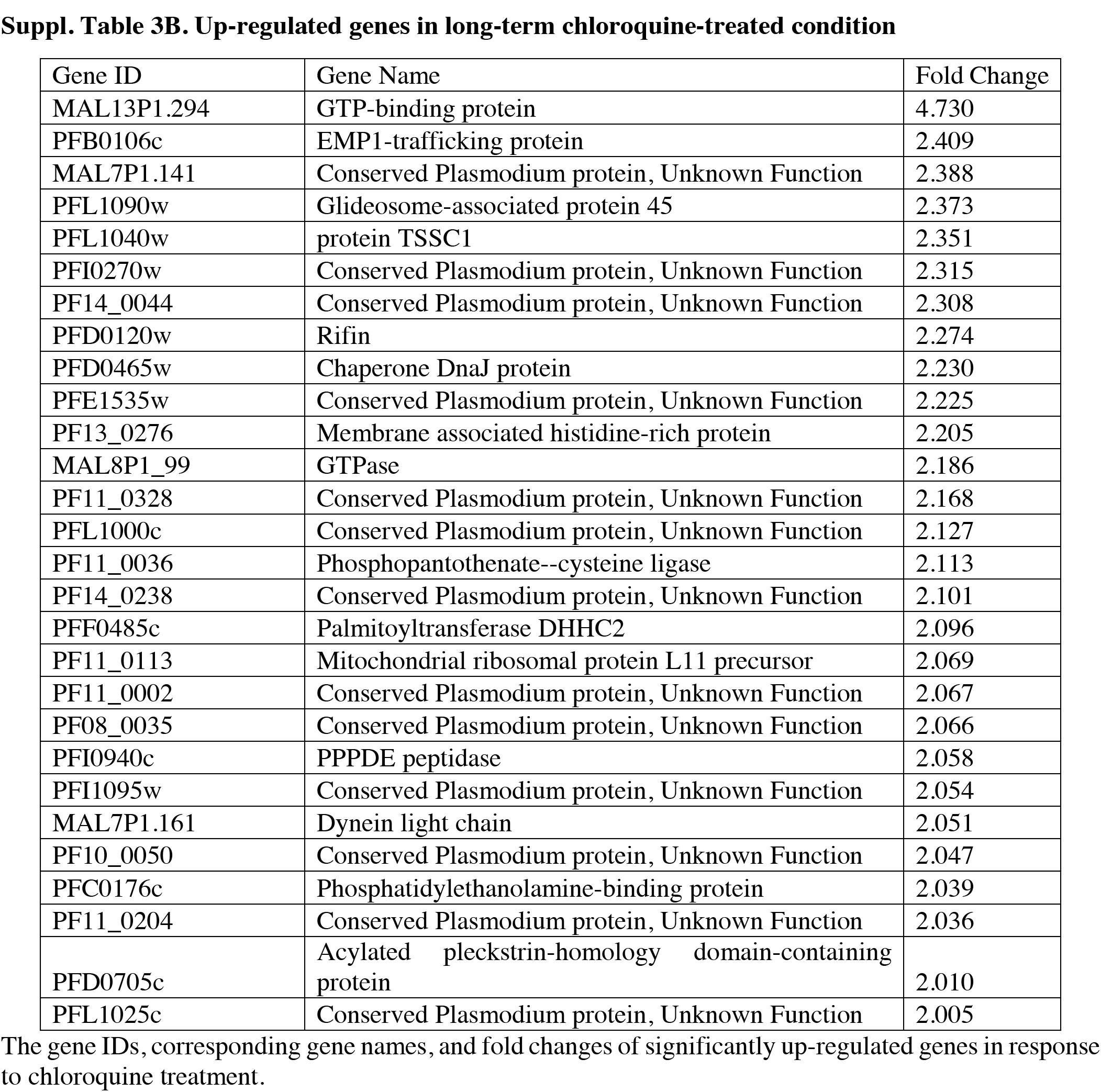
Up-regulated Differentially Expressed Genes in (A) Short and (B) Long-term Chloroquine-treated Parasites.

**Suppl. Table 4:**
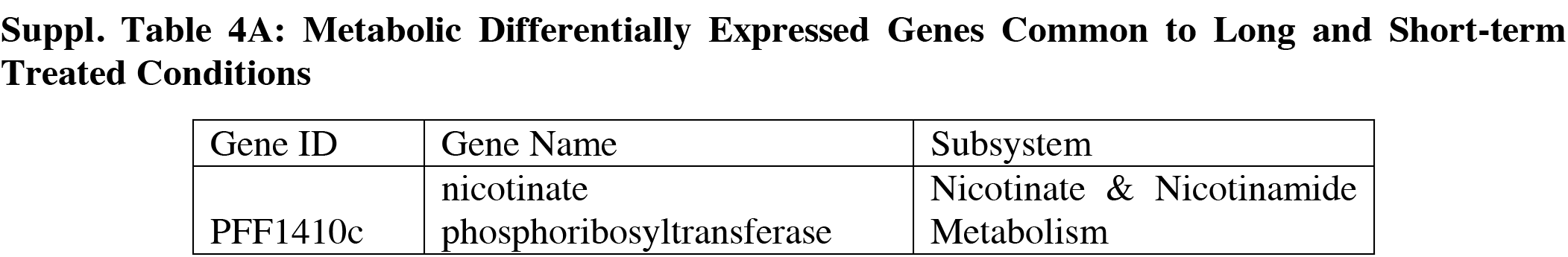
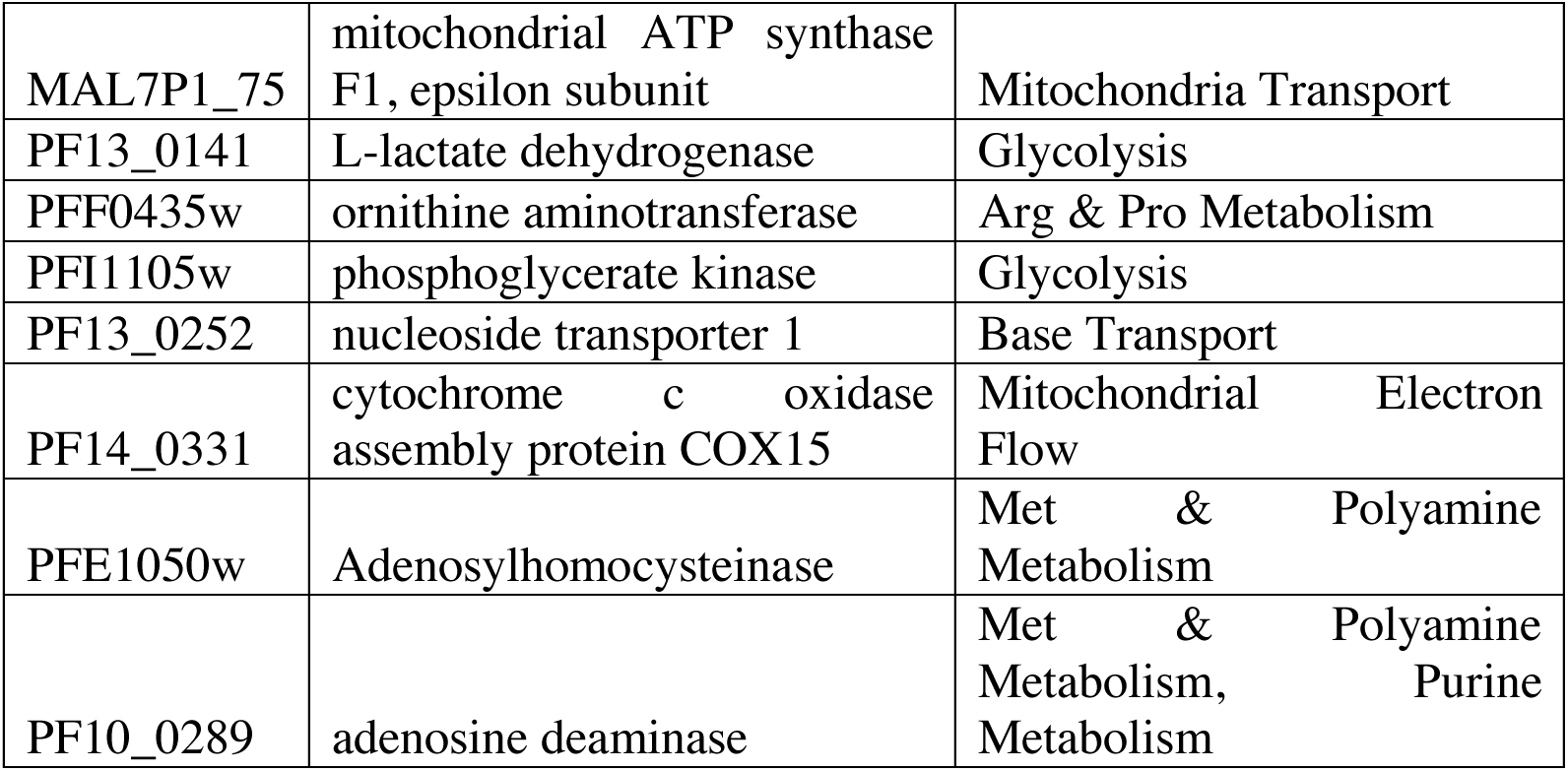
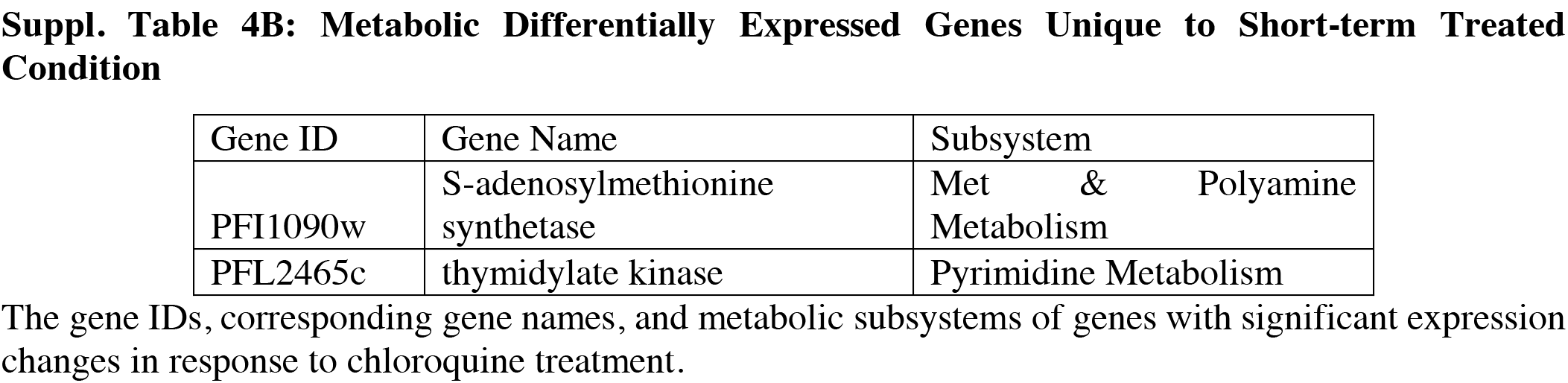
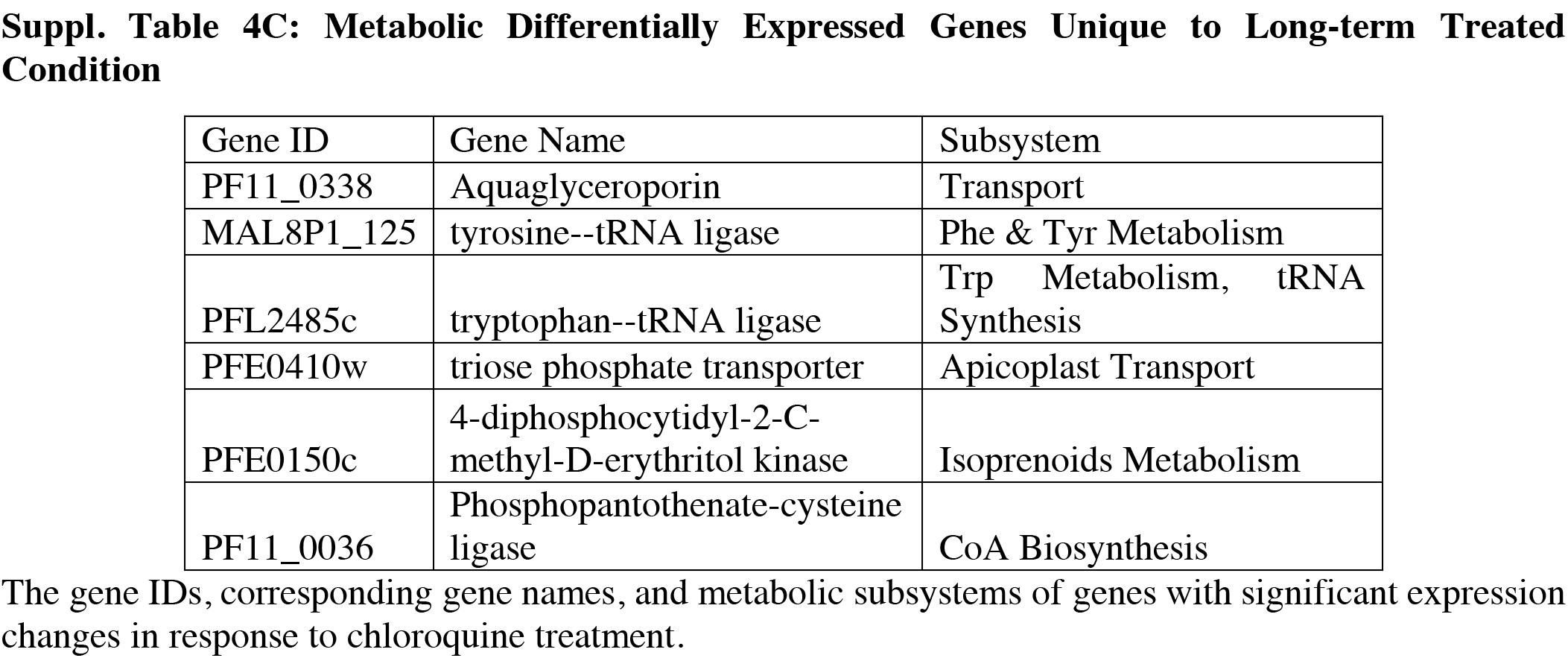
Significant Down Regulation of Metabolic Genes Arising from Chloroquine Treatment.

**Suppl. Table 5.**
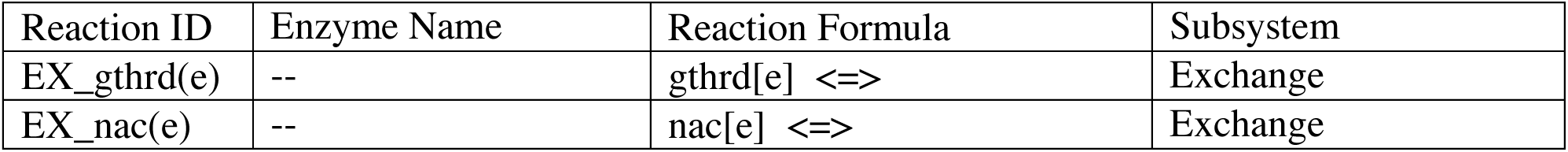
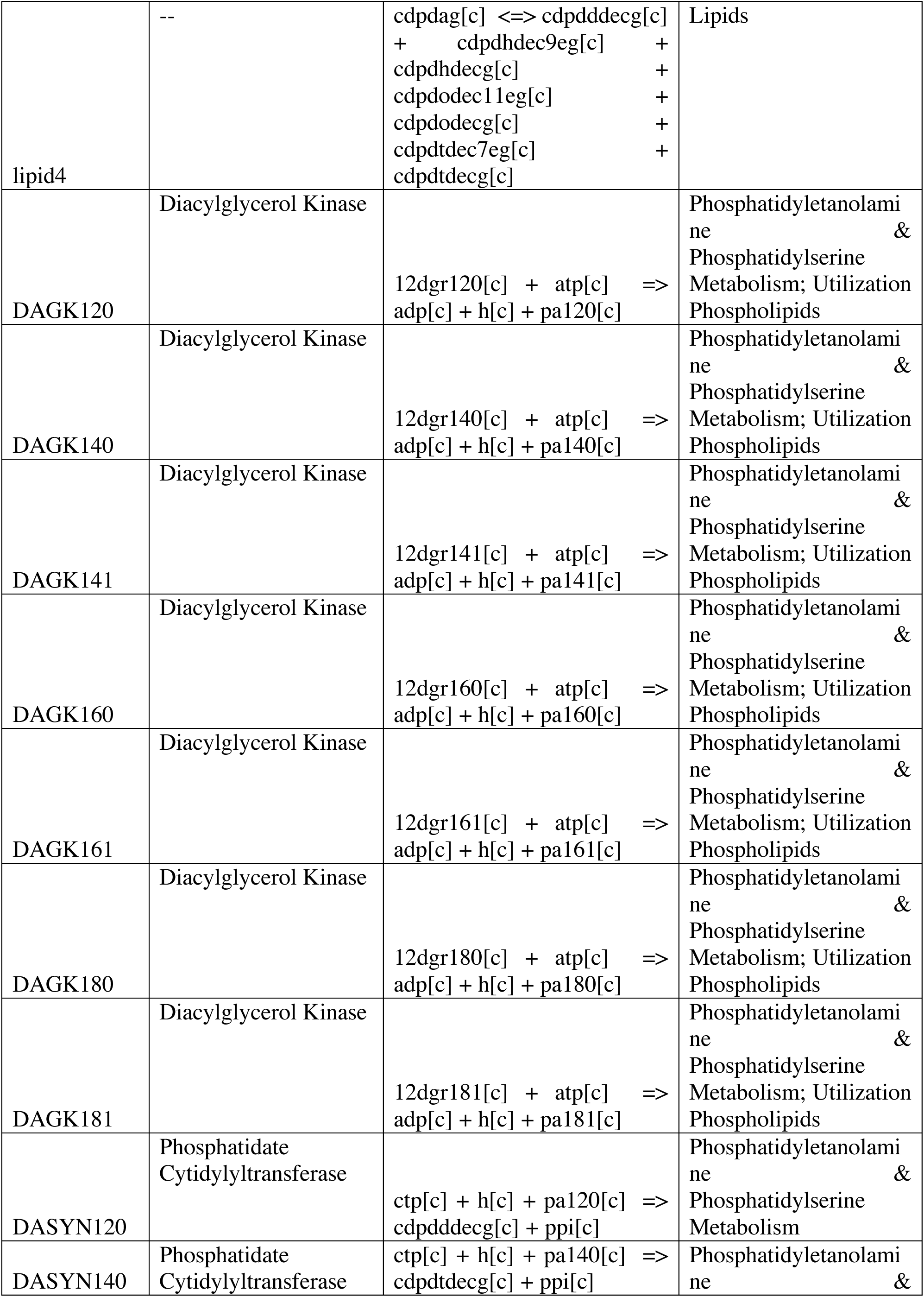
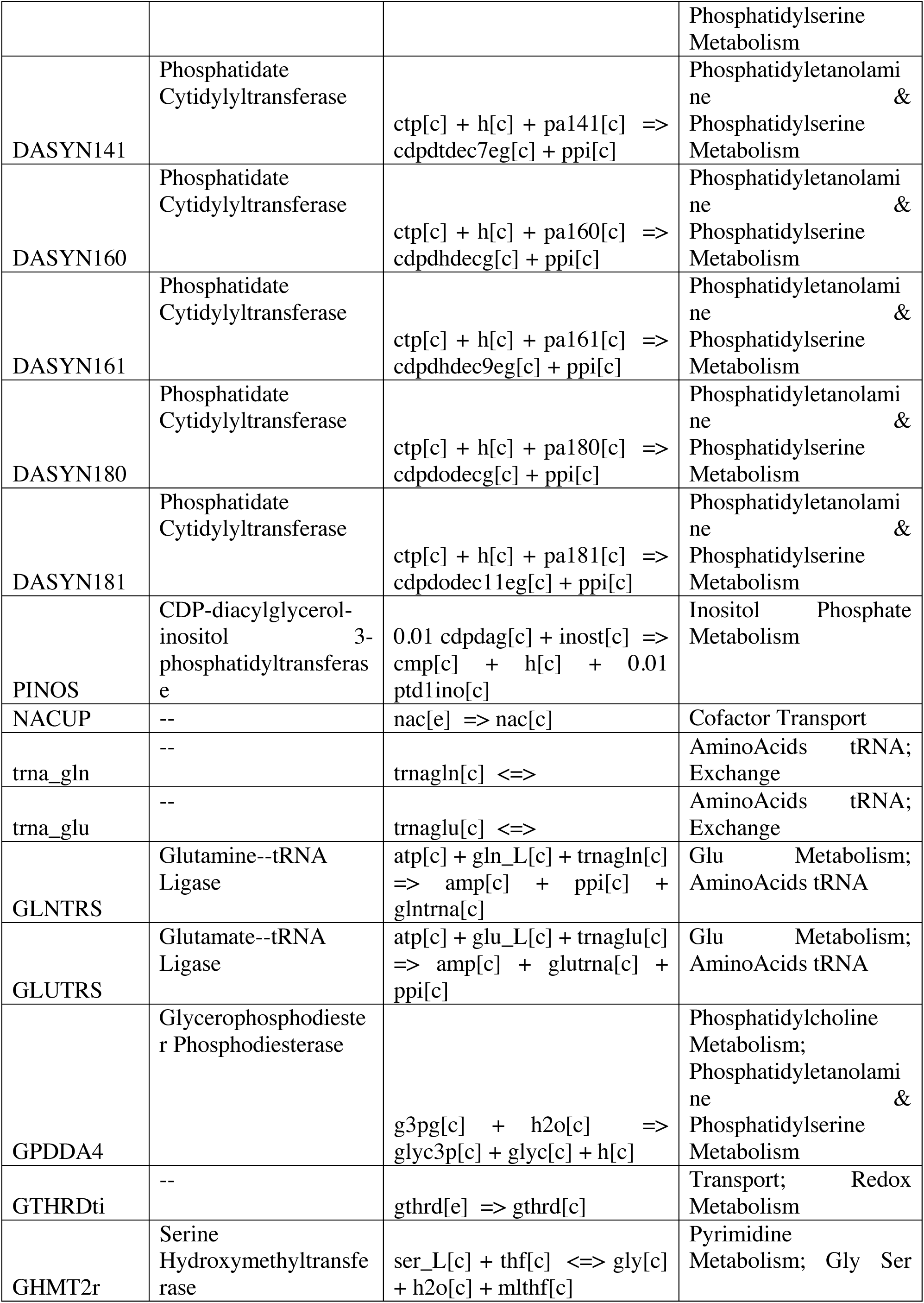
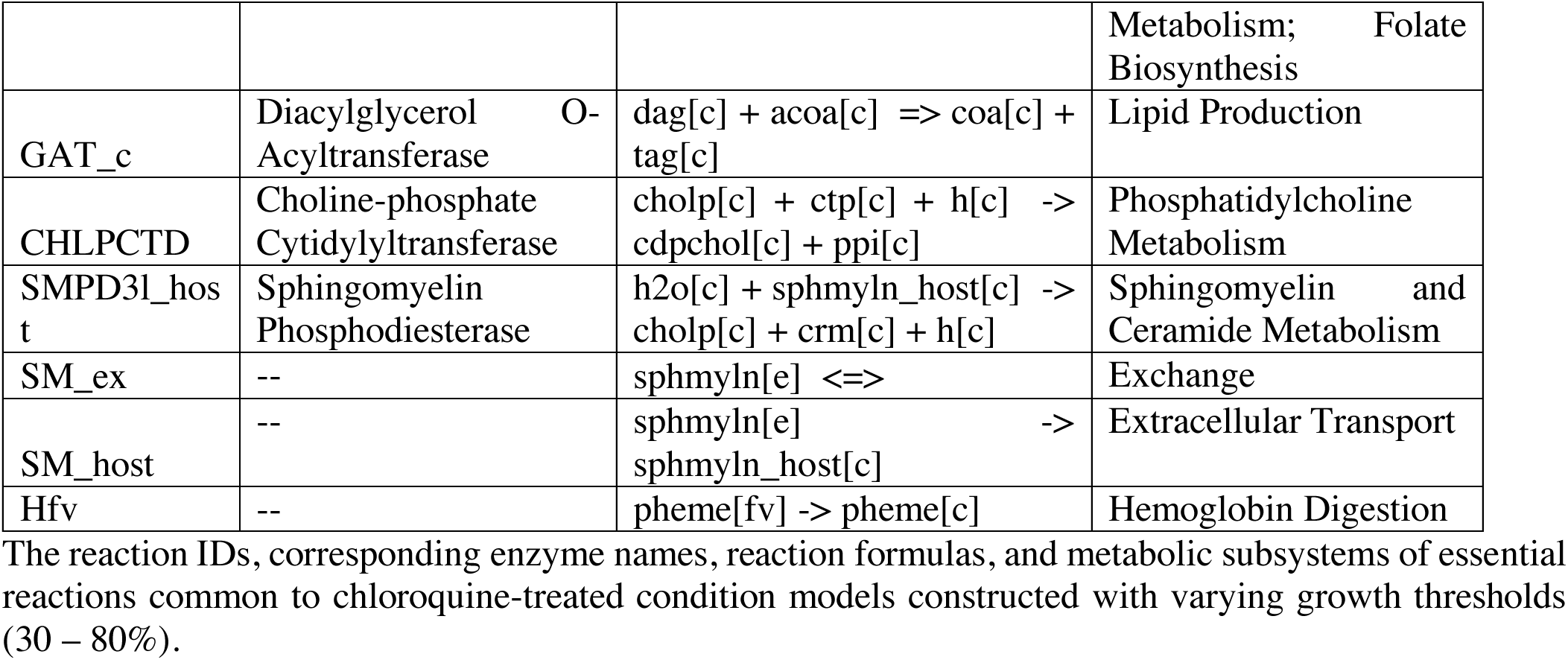
Reactions Essential in only Chloroquine-Treated Models.

**Suppl. Table 6.**
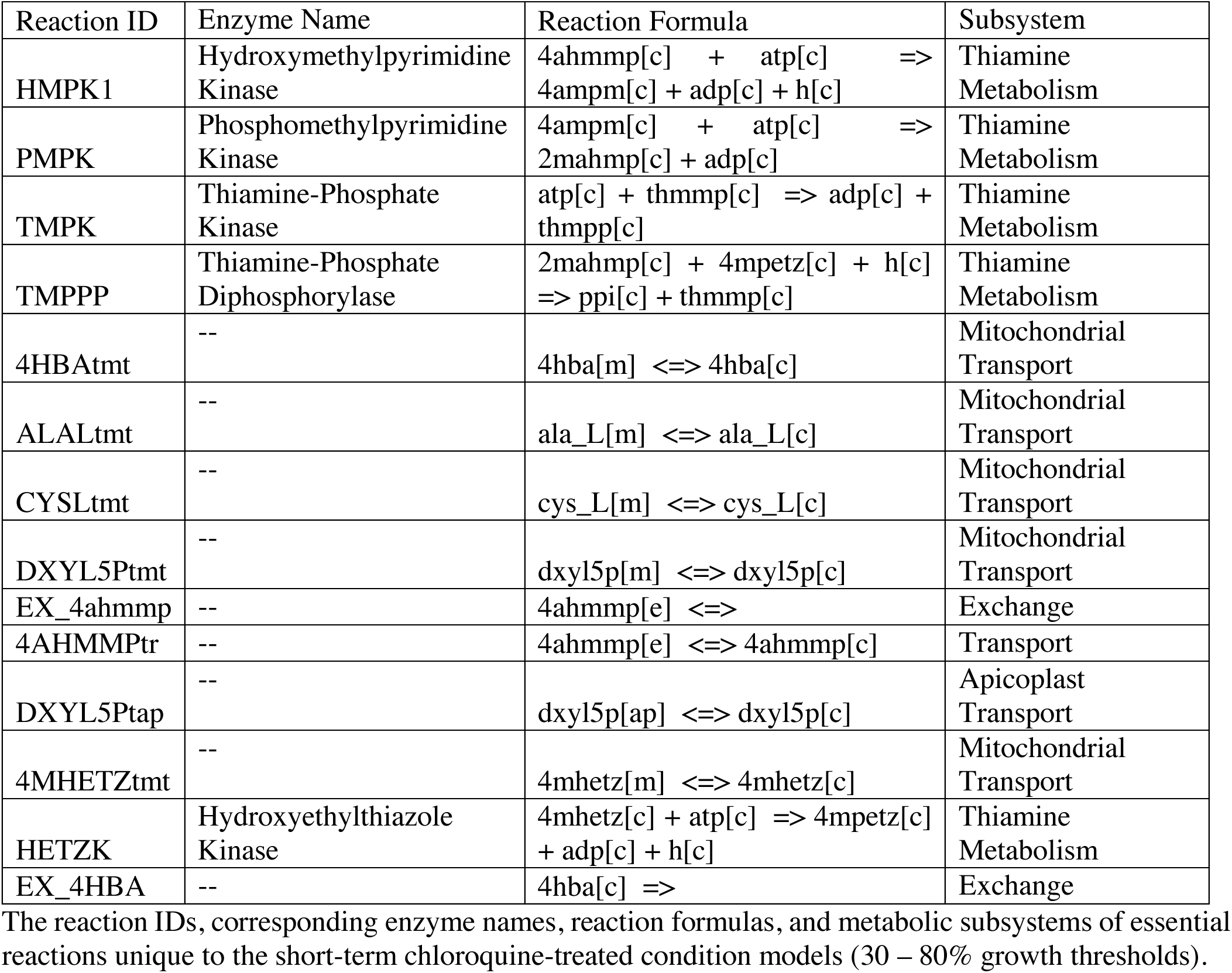
Reactions Essential in only the Short-Term Treatment Condition.

**Suppl. Table 7.** Reactions Essential in only the Long-Term Treatment Condition.

| Reaction ID | Enzyme | Reaction Formula | Subsystem |
| --- | --- | --- | --- |
| PGMT_2 | Phosphoglucomutase | $r1p[c] \rightleftharpoons r5p[c]$ | Pentose Phosphate Cycle; Glycolysis |
| ADPtap | -- | $adp[ap] \rightleftharpoons adp[c]$ | Apicoplast Transport |
| ATPtap | -- | $atp[ap] \rightleftharpoons atp[c]$ | Apicoplast Transport |
| PYK | Pyruvate Kinase | $adp[c] + h[c] + pep[c] \Rightarrow atp[c] + pyr[c]$ | Glycolysis; Pyruvate Metabolism; Isoprenoids Metabolism; Fatty Acid Synthesis |
| ADCL | Aminodeoxychorismate Lyase | $4adcho[c] \Rightarrow 4abz[c] + h[c] + pyr[c]$ | Shikimate Biosynthesis; Folate Biosynthesis |
| ADCS | para-Aminobenzoic Acid Synthetase | $chor[c] + gln\_L[c] \Rightarrow 4adcho[c] + glu\_L[c]$ | Folate Biosynthesis; Shikimate Biosynthesis |

3 The reaction IDs, corresponding enzyme names, reaction formulas, and metabolic subsystems of essential 4 reactions unique to the long-term chloroquine-treated condition models (30 – 80% growth thresholds).

**Suppl. Table 8.**
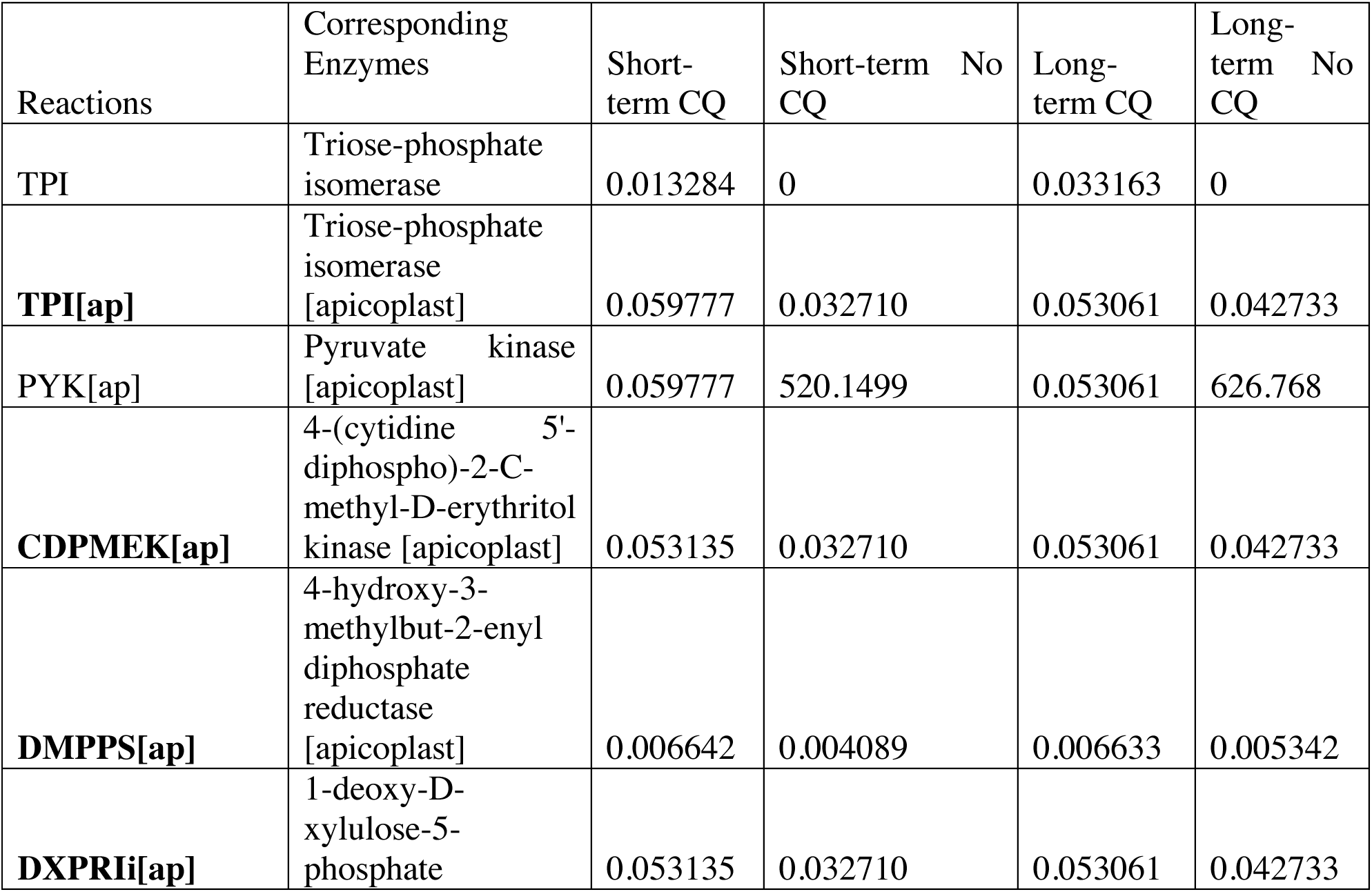
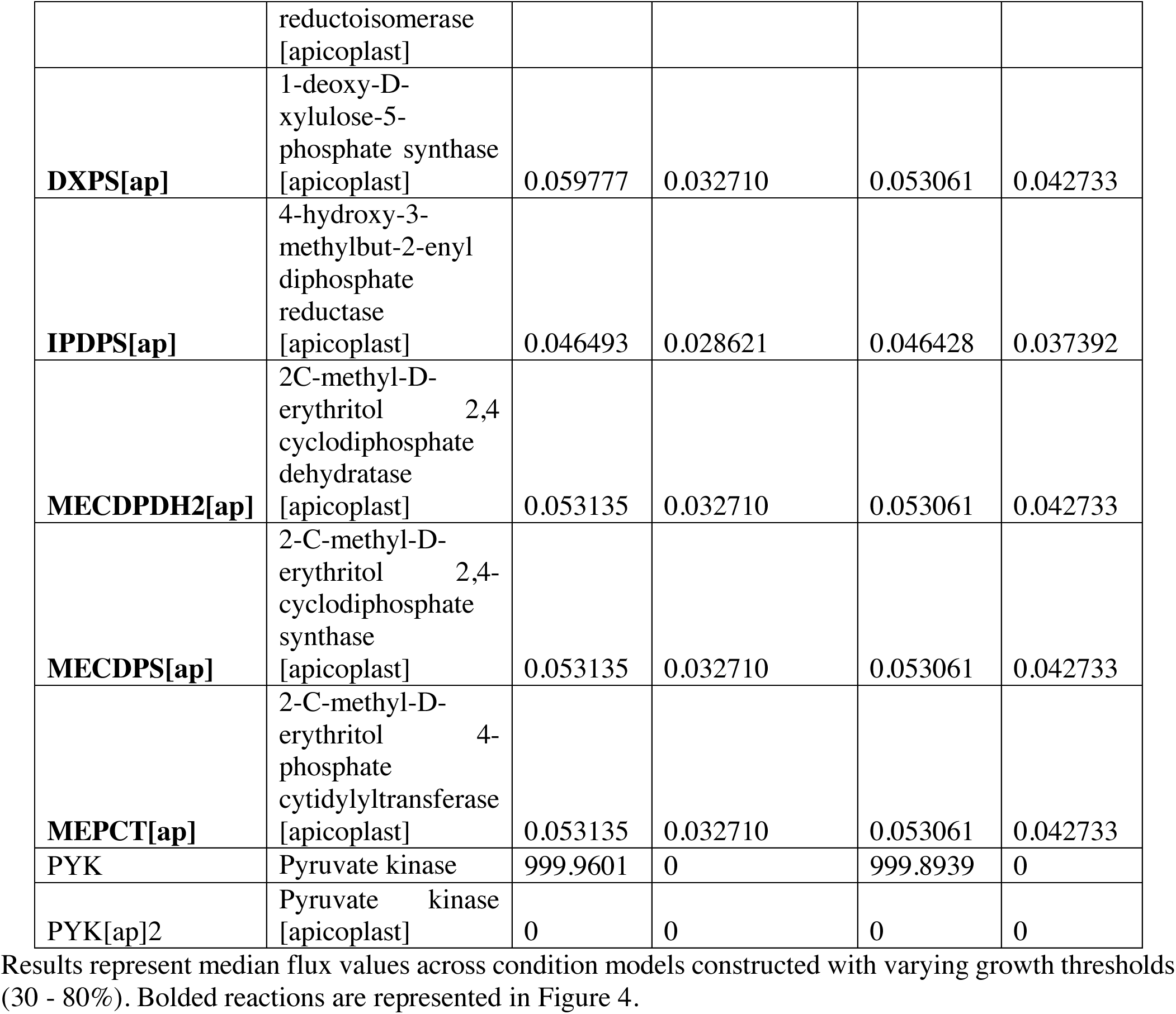
Flux Values for Isoprenoid Metabolism Reactions in the Chloroquine-treated and Untreated Models.

**Suppl. Table 9.** Flux Values for Folate Metabolism Reactions in the Chloroquine-treated and Untreated Models.

| Reactions | Corresponding Enzymes | Short-term CQ | Short-term No CQ | Long-term CQ | Long-term No CQ |
| --- | --- | --- | --- | --- | --- |
| **FMETTRS** | Methionyl-tRNA<br>formyltransferase | 0.297843 | 0.183353 | 0.297428 | 0.239536 |
| AKP1 | Alkaline phosphatase | 0.059777 | 0.032710 | 0.053061 | 0.042733 |
| **DHFR** | Dihydrofolate reductase | 0.830357 | 0.689333 | 0.829198 | 0.667803 |
| **DHFS** | Dihydrofolate synthase | 0.019926 | 0.016542 | 0.019898 | 0.016025 |
| **DHPS2** | Dihydropteroate synthase | 0.019926 | 0.016542 | 0.019898 | 0.016025 |
| GTPCI | GTP cyclohydrolase I | 0 | 0 | 0 | 0 |
| **HPPK2** | 2-amino-4-hydroxy-6-hydroxymethyldihydropteridine<br>diphosphokinase | 0.019926 | 0.016542 | 0.019898 | 0.016025 |
| **MTHFC** | Methenyltetrahydrofolate cyclohydrolase | 0.304485 | 0.252773 | 0.30406 | 0.244878 |
| **MTHFD** | Methylenetetrahydrofolate dehydrogenase (NADP+) | 0.304485 | 0.252773 | 0.30406 | 0.244878 |
| PTHPS | 6-pyruvoyltetrahydropterin synthase | 0 | 0 | 0 | 0 |
| THFGLUS | Tetrahydrofolate synthase | 0 | 0 | 0 | 0 |
| **TMDS** | Thymidylate synthase | 0.810432 | 0.672792 | 0.8093 | 0.651778 |
| PTHPS2 | 6-pyruvoyltetrahydropterin synthase | 0 | 0 | 0 | 0 |
| **ADCL** | 4-aminobenzoate synthase | 0 | 0 | 0.019898 | 0 |
| **ADCS** | 4-amino-4-deoxychorismate synthase | 0 | 0 | 0.019898 | 0 |
| **GHMT2r** | Glycine hydroxymethyltransferase | 1.121559 | 0.931078 | 1.119993 | 0 |
| GHMT2r_m | Glycine hydroxymethyltransferase [mitochondria] | 0 | 0 | 0 | 0 |
| GHMT2r_ap | Glycine hydroxymethyltransferase [apicoplast] | 0 | 0 | 0 | 0 |
| **pABAt** | Folate transporter 1 & 2 | 0.019926 | 0.012266 | 0 | 0.016025 |

## Ethics approval and consent to participate

Not applicable.

## Consent for publication

Not applicable.

## Availability of data and materials

The publically available dataset analyzed during the current study are available on the NCBI’s Gene Expression Omnibus, GSE31109. Additionally, our curated model and code are available on GitHub, at https://github.com/anauntaroiu/Chloroquine-Project.

## Authors’ contributions

AU and MC designed the study. AU curated the model and performed statistical and network analyses. AU and MC interpreted the data and analyses. AU wrote the manuscript. MC, JG, and JP edited the manuscript. JG and JP provided resources. All authors read and approved the final manuscript.

## Acknowledgements

We thank the members of the Guler and Papin labs for their thoughtful feedback and the openCobra community for software development integral to this work.

## List of abbreviations

CQ: chloroquine
DEG: differentially expressed gene
FBA: flux balance analysis
FC: fold change
FDR: false-discovery rate
MADE: Metabolic Adjustment for Differential Expression

